# NEST: Spatially-mapped cell-cell communication patterns using a deep learning-based attention mechanism

**DOI:** 10.1101/2024.03.19.585796

**Authors:** Fatema Tuz Zohora, Eugenia Flores-Figueroa, Joshua Li, Deisha Paliwal, Faiyaz Notta, Gregory W. Schwartz

**Affiliations:** Princess Margaret Cancer Centre, University Health Network, Toronto, ON M5G 1L7, Canada; Department of Medical Biophysics, University of Toronto, Toronto, ON M5G 1L7, Canada; Vector Institute for Artificial Intelligence, 661 University Ave Suite 710, Toronto, ON M5G 1M1; PanCuRx Translational Research Initiative, Ontario Institute for Cancer Research, Toronto, ON, M5G 0A3, Canada

## Abstract

Dysregulation of communication between cells mediates complex diseases such as cancer and diabetes. However, detecting cell-cell communication (CCC) at scale remains one of the greatest challenges in transcriptomics. While gene expression measured with single-cell RNA sequencing and spatial transcriptomics reinvigorated computational approaches to detecting CCC, most existing methods exhibit high false positive rates, do not integrate spatial proximity of ligand-receptor interactions, and cannot detect CCC between individual cells. We overcome these challenges by presenting *NEST (NEural network on Spatial Transcriptomics)*, which uses a graph attention network paired with an unsupervised contrastive learning approach to decipher patterns of communication while retaining the strength of each signal. We introduce new synthetic benchmarking experiments which demonstrate how NEST outperforms existing tools and detects biologically-relevant CCC along with directionality and confidence across spot- and cell-based technologies measuring several different tissues and diseases. In our applications, NEST identifies T-cell homing signals in human lymph nodes, aggressive cancer CCC in lung adenocarcinoma, and discovers new patterns of communication that act as relay networks in pancreatic cancer. Beyond two-dimensional data, we also highlight NEST’s ability to detect CCC in three-dimensional spatial transcriptomic data.

## Introduction

Cell-cell communication (CCC) enables the complex coordination of cells, forming tissues and organs in multicellular organisms and accomplishing critical biological functions. How-ever, aberrant communication among cells or atypical decoding of molecular messages can lead to and promote diseases such as cancer. CCC is involved in several hallmarks of cancer, such as tumor-promoting inflammation, inducing or accessing vasculature, and activating invasion and metastasis^1–4^. It is therefore crucial to pinpoint communication responsible for normal and aberrant cell and tissue function to inform the next generation of therapeutics.

CCC is mediated by ligand-receptor pairs where a “sender” cell produces ligand proteins which bind to matching receptor molecules on a “receiver” cell^4^. Common techniques to identify CCC use single-cell RNA sequencing (scRNA-seq) data by matching highly expressed ligand genes from the sender cell type with highly expressed receptor genes from a receiver cell type, prioritizing ligand-receptor pairs with high “ligand-receptor co-expression scores”. These scores represent the overall expression of the ligand-receptor pair. After identifying ligand-receptor pairs, these tools diverge by determining confidence in each pair using statistical tests^5–7^, substituting receptor genes with pathways^8^, or using graph-based approaches^9^. Others, like CellChat^10^, use network analysis and pattern recognition approaches. NicheNet^11^ uses signaling pathway networks and the PageRank algorithm. Despite advances proposed by these methods, detecting CCC remains a major challenge.

Past efforts to identify CCC have high false positive and negative rates^12^, which is in part due to using a single data modality — the transcriptome — from cells. Only 6% of genes exhibit significant expression changes in response to ligands, which may contribute to low accuracy without additional context such as neighboring cells^13^. Furthermore, scRNA-seq requires tissue dissociation, destroying the spatial context of cells. As CCC is spatially dependent, scRNA-seq introduces challenges for true single-cell CCC detection instead of cell-type communication^14^. Lately, methods such as Scriabin^15^ and GraphComm^16^ detect CCC from scRNA-seq data alone and map final results to spatial regions within tissue using corresponding spatial transcriptomic data. However, these approaches incorporate spatial position not to detect CCC, but to validate already found CCC among adjacent cells. Such methods also do not report distant ligand-receptor interactions such as paracrine interactions, which constitute the majority of most ligand-receptor databases. To overcome such limitations, new CCC models directly integrating the spatial context of gene expression are necessary.

Spatial transcriptomic technologies such as Visium^17,18^ and multiplexed error-robust fluorescence in situ hybridization (MERFISH)^19^ measure the physical location of cells paired with their transcripts, providing new opportunities to detect CCC. Visium measures transcriptomes of barcoded spots of 55 µm in diameter, each containing about 1-10 cells, while the recent launch of Visium HD (high definition) achieves single-cell spatial resolution at 2 µm. Alternatively, MERFISH provides true single-cell resolution of individual cells albeit with a smaller subset of genes. There are few existing methods which detect CCC directly from spatial transcriptomic data, informed by both the spatial location of cells and their associated transcripts. Niches^20^ uses k-nearest neighbors to identify proximal cells and calculates their ligand-receptor co-expression scores. Niches then collapses cells to neighborhoods using principle component analysis (PCA) to discover niches of communication. COMMOT^21^ screens cell–cell communication in spatial transcriptomics via collective optimal transport. However, COMMOT requires a network pathway list as additional input, which increases its reliance on *a priori* information. Most of these methods use differentially expressed and variable ligand and receptor genes, only incorporating spatial information to limit potential communication to a neighborhood of cells. However, spatial data can offer additional information such as discovering re-occuring CCC “patterns”. These patterns expand CCC definitions beyond ligand-receptor pairs to ligand-receptor-ligand-receptor relay networks (ligand from one cell to a receptor on another cell which sends another ligand to a third cell’s receptor, and so on). The frequency of these patterns may indicate higher confidence in CCC detection^22^. Discovery of such CCC patterns requires sophisticated artificial intelligence models to learn complex structures.

To facilitate CCC detection, we can represent communication from spatial transcriptomic data as a knowledge-graph where cells or spots are vertices and edges represent different types of neighborhood relations. As our goal is to predict which relations are probable communication, a deep learning option to unravel the communication network is a graph neural network (GNN)^23^. GNN is an effective model for encoding topological structures in graph representations by generating a graph embedding. Variants of GNN are already being applied in transcriptomic data, including a graph convolution network for clustering^24^ and a GNN-based encoder for deconvolution and integration^25^. A newer addition to the transformer^26^ family is the graph attention network (GAT), a powerful tool that has already revolutionized other knowledge-graph-based problems such as social networks^27^ and molecular structures^28^. As this model requires ground truth data for supervised model training, we propose using a contrastive learning approach, Deep Graph Infomax (DGI)^29^, which excels in unsupervised learning problems^24,30,31^.

Built with these state-of-the-art advances in artificial intelligence, we present *NEST (NEural network on Spatial Transcriptomics)*, a method to measure cell-cell communication and patterns between individual cells that leverages a GAT encoder model with DGI contrastive learning. Using new benchmarks for single-cell ligand-pair detection and CCC patterns, we demonstrate that NEST outperforms existing tools. We applied our model to four data sets across multiple tissues, species, and technologies^32–34^ to map spatially-resolved CCC. Importantly, we show that NEST can accurately reconstruct not only traditional single ligand-receptor signals between cells, but also relay network patterns of communication in both 2D and 3D spatial transcriptomic samples. Applying NEST to our cohort of pancreatic ductal adenocarcinoma (PDAC) patients revealed critical CCC associated with PDAC progression and spatially associated with known PDAC subtypes linked with treatment response and overall survival. As demonstrated, NEST is not limited to a single technology or species. Rather, it is a transferable model applicable to data across domains. We believe NEST is a major step forward in accelerating the application of deep learning in spatial transcriptomics as well as other related knowledge-graph-based contexts. NEST is open-source and publicly available at https://github.com/schwartzlab-methods/NEST.

## Results

### NEST identifies patterns of communication in spatial transcriptomic data

Ligand-receptor pair-based communication depends on spatial distance; however, the vast majority of existing tools do not leverage positional information to detect CCC and collapse communicating units to cell types and clusters rather than spots and cells. To overcome these limitations, we developed NEST for high-resolution, spatially-resolved CCC detection (Figure 1).

**Figure 1:**
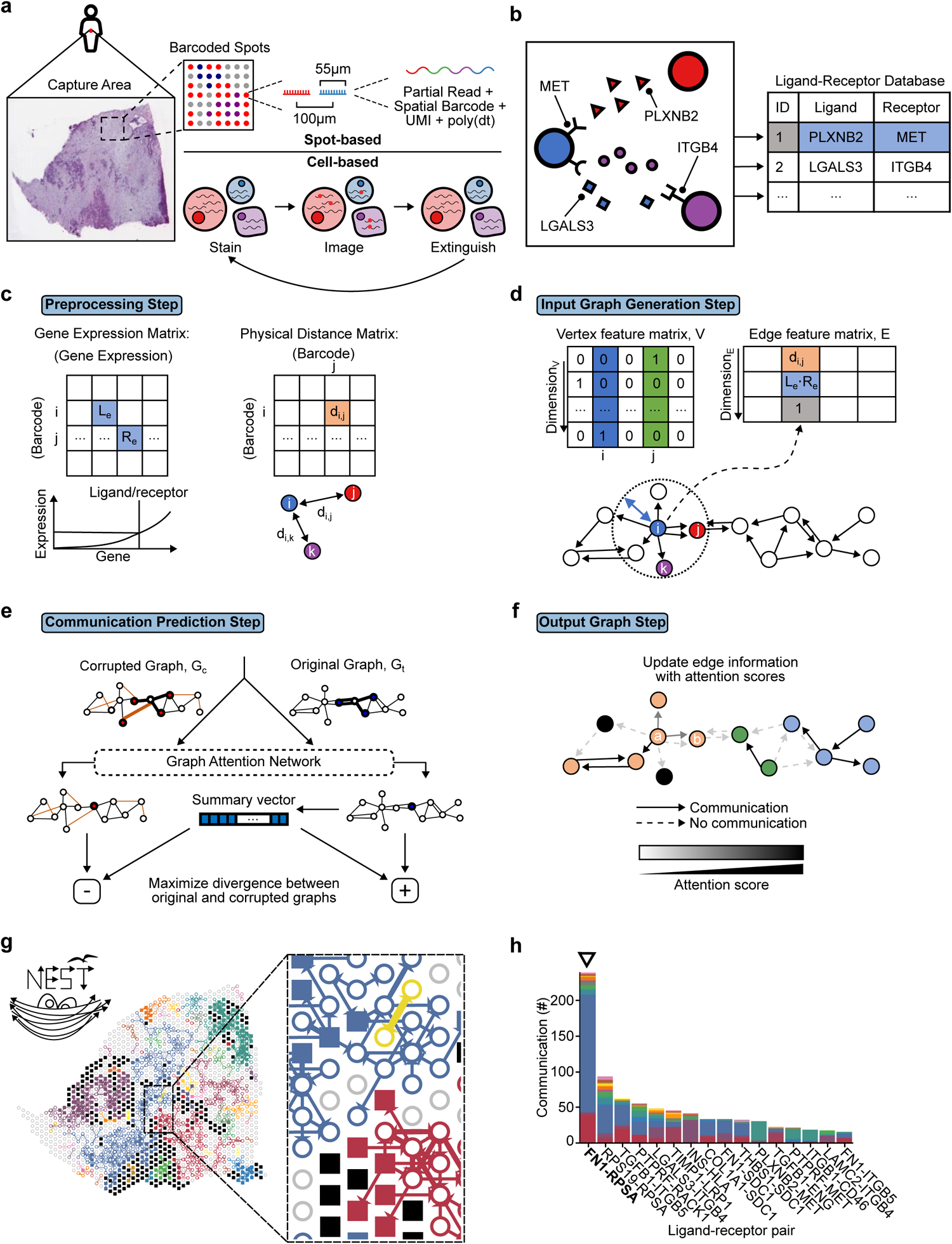
Overview of detecting cell-cell communication with NEST. **a** Input tissue sample at either spot (e.g. Visium; top) or cell resolution (e.g. MERFISH; bottom) data format. **b** Input ligand-receptor database containing known ligand and cognate receptor pairings. **c** Preprocessing Step, where genes with an expression above a threshold percentile are considered active (left). Pairwise Euclidean distance between vertices are stored in a physical distance matrix (right). **d** Input Graph *G* = (*V*, *E*) Generation Step with *V* spots or cells as vertices and *E* edges as neighborhood relations, some of which represent communication (bottom). An input threshold distance is used for the neighborhood formation (blue arrow). From the graph, vertex features are represented as a one-hot vector matrix (top left). The edge feature matrix holds edge feature vectors containing three attributes: pairwise distance, ligand-receptor co-expression score, and the ligand-receptor pair identity from the database in **b**. **e** Communication Prediction Step using a GAT encoder through unsupervised contrastive learning with DGI. **f** Output Graph Step visualizing edges with the highest attention scores. Attention scores range from 0 (white) to 1 (black), where 1 represents the strongest connections. Lower-scoring edges are removed (dashed lines), resulting in subgraphs of communicating vertices. **g** Example output showing the flow of communication between tumor-annotated spots (filled squares) with stroma spots (open circles), colored by connected component. **h** An example NEST-generated histogram showing the frequency of communication through ligand-receptor pairs in those top 20% highly attention scored edges. Colors in the histogram correspond to the colors of subgraphs in **g**. For instance, the most abundant communication labeled as *FN1*-*RPSA* is found primarily in the blue region. Together, NEST presents a high-resolution solution to detect the strength and location of cell-cell communication on tissue.

Given a 2D or 3D spatial transcriptomic data set at either spot or single-cell resolution and an existing ligand-receptor database, NEST scores each intercellular signal based on the co-expression of highly-expressed ligand and cognate receptor genes (Figure 1a-c). To achieve true single-cell and -spot level communication identification, NEST relies on a GNN, a class of deep-learning-based models, to identify which ligand-receptor pairs are highly probable to exist in a tissue sample based on re-occurring patterns of communication in a particular region. For example, TGF*β*1 signaling is upregulated in tumor cells across various cancers^33^. This signal occurs multiple times in cancer tissue along the boundary of tumor and non-tumor cells, forming a distinct pattern that is not observed in other regions of the same tissue. Patterns may be more than a single ligand-receptor pair, however, as there may be CCC acting as a relay network with multiple “hops” between cells^22^. Deep learning models excel in detecting such hidden patterns, so NEST leverages this power by using a graph attention network^35,36^ (GAT), an encoder model that records such patterns in the form of a vertex embedding.

After data preprocessing, NEST converts spatial transcriptomic data into a graph *G* = (*V*, *E*) with *V* cells or spots as vertices and *E* edges as some neighborhood relation between the pair of vertices (Figure 1d; see Methods). NEST inserts edges between a pair of cells or spots (herein referred to as vertices) *i* and *j* if they are proximal neighbors with *i* expressing high ligand and *j* expressing high receptor (high co-expression). *G* can be massive, containing thousands of vertices with millions of edges based on the number of expressed genes. Importantly, *E* represents neighborhood relations and not CCC, as proximal cells do not always establish communication. Tissue context^13^, epigenetic factors^37^, and other signaling pathways^11^ may influence high ligand-receptor co-expression. NEST sifts through these putative relations to decide which represents real communication. For this purpose, we pass *G* to the core deep learning module in NEST, the “Communication Prediction” step, where a GAT model generates the vertex embedding (Figure 1e).

The traditional GAT model requires ground truth data for training an encoder, but this information is unknown from spatial transcriptomic data. We instead chose to implement unsupervised training through DGI^29^, a contrastive learning approach. NEST, to our knowledge, is the first method of its kind to apply DGI to GATs (Figure 1e and Supplementary Figure S1a). DGI compares encoder weights derived from the observed network with encoder weights from a “corrupted” network of randomly shuffled and permuted vertices and edges. DGI maximizes weights from the observed network while penalizing weights from the corrupted network. As the model converges, NEST assigns higher attention scores to stronger neighborhood relations (Figure 1f). We use these attention scores to represent communication strength. To retain the most probable intercellular signals, we filter edges, keeping the top 20% of highly attention-scored edges by default.

After detecting high-resolution CCC, NEST also identifies highly communicating regions of tissue in the “Output Graph” step by determining connected components^38^ (Figure 1f, g). As NEST identifies CCC between each vertex along with signal strength, we provide a unique visualization that displays vertices colored by densely communicating regions of tissue along with ligand-receptor pairs as an arrows whose thicknesses are determined by their attention scores (Figure 1g). To complement the tissue visualization, NEST also generates histograms showing the count of all ligand-receptor pairs in the top-most highly attention-scored edges colored by the community they are found in within the tissue (Figure 1h). With this extensive tool set, NEST is fully equipped as an end-to-end framework for spatially-resolved CCC detection.

### NEST pinpoints T-cell homing signals in T-cell zones of the human lymph node

To determine the accuracy of our algorithm, we applied NEST to Visium data from a human lymph node (Figure 2a-e) with *n* = 4,035 spots^32^. We hypothesized that NEST would identify the T-cell homing signal of chemokine (C–C motif) ligand 19 with cognate CC-chemokine receptor 7 (*CCL19*-*CCR7*) and place this CCC within the T-cell zone^39,40^. The T-cell zone was previously annotated using cell2location^32^ (Figure 2a). We applied NEST to the entire tissue and ranked all CCC based on their attention scores, keeping the ligand-receptor pairs with the top 20% highest attention scores and located within the T-cell zone (Figure 2b). Among the 12,605 possible ligand-receptor pairs in the database, NEST identified *CCL19*-*CCR7* as the second most abundant pair in the T-cell zone, with strict thresholds above 20%. The topmost detected pair was *CCL21*-*CXCR4*, another T-cell migratory signal^41^ (Figure 2b). Surprisingly, while NEST found *CCL19*-*CCR7* as a top signal in the T-cell zone based on attention score, this pair’s co-expression score was not among the highest (Figure 2c,d). Increasing attention score thresholds further confirmed T-cell zones as the primary location for *CCL19*-*CCR7* (Figure 2e). This prioritization and localization suggest that NEST does not score edges based solely on input ligand and receptor expression. Instead, NEST focuses on hidden communication patterns to decide which edges are essential to represent the context of the tissue sample.

**Figure 2:**
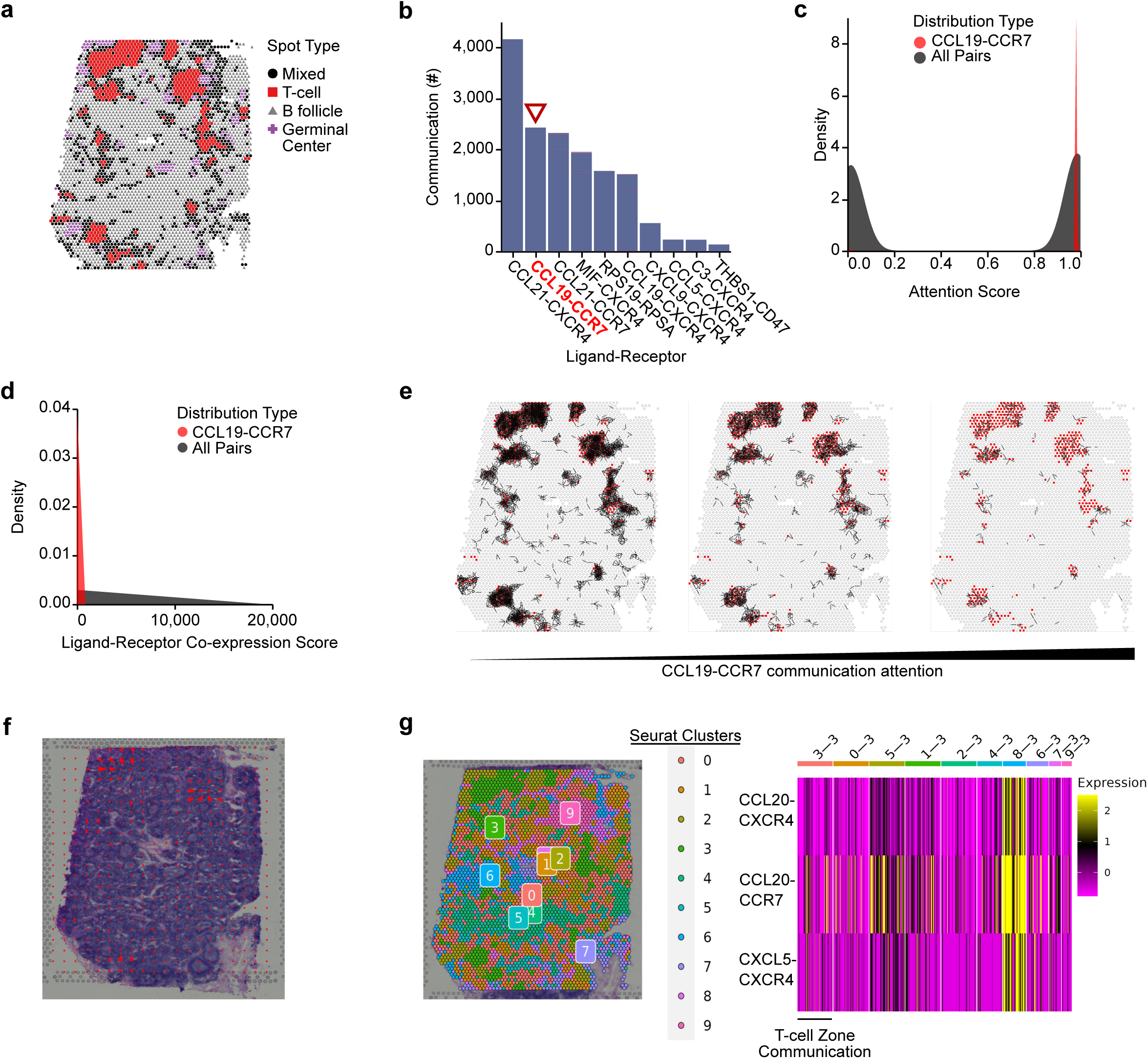
NEST identifies T-cell homing signals in human lymph node T-cell zones. **a** Human lymph node tissue measured with Visium^32^. **b** Histogram of ligand-receptor pairs (*x*-axis) with the top 20% highest attention scores in the T-cell zone assigned by NEST, in descending order of abundance (*y*-axis). *CCL19*-*CCR7* (red text with triangle) is a canonical T-cell homing signal. **c** Density plot of *CCL19*-*CCR7* attention scores (red) compared to all other ligand-receptor pairs (gray) in T-cell zone. **d** Density plot of *CCL19*-*CCR7* ligand-receptor co-expression scores (red) compared to all other ligand-receptor pairs (gray) in T-cell zone. **e** Selection of the top 5000, 2500, and 500 *CCL19*-*CCR7* edges with the strongest attention scores (left to right) across the entire tissue. Stronger *CCL19*-*CCR7* communication is found in the T-cell zone. **f** Application of COMMOT to the human lymph node, with red arrows indicating *CCL19*-*CCR7* strength. **g** Application of Niches to the human lymph node. Using a cluster-based analysis (left), Niches identifies three signals excluding *CCL19*-*CCR7* signal (right).

To compare the performance of NEST against other emerging tools for CCC detection, we applied Niches and COMMOT to identify *CCL19*-*CCR7* within the T-cell zone. COMMOT, while detecting *CCL19*-*CCR7* in the largest major T-cell zone, was unable to capture the signal across the whole tissue even with the additional requirement of a communication pathway data set (Figure 2f). Niches detected three signals in total within T-cell zones, but none are as highly referenced as the canonical *CCL19*-*CCR7* (Figure 2g). Based on this comparative analysis of our model and other existing tools, NEST alone can detect CCC in specific regions of tissue accurately and may complement COMMOT and Niches for spatial transcriptomic analysis.

### NEST improves communication detection in robust benchmarks across cell- and spot-based distributions

Confirming that NEST had superior quantitative and visual detection of biologically relevant CCC in the lymph node samples, we sought to quantify such results with robust bench-marking. As our benchmarks required ground truth data which is unavailable for biological samples, we prepared 12 synthetic data setups across three spatial distributions to quantify NEST’s performance (Figure 3). We simulated gene expression and coordinates to represent spots or individual cells, analogous to Visium and MERFISH technologies, respectively. We introduced communication by assigning higher gene expression to genes representing synthetic ligand-receptor pairs in randomly selected data points. We evaluated each method on whether the model can detect the sender, receiver, and exact ligand-receptor pair. To represent different distributions of cells and spots, we compared methods across three types of benchmarks: equidistant data points (*n* = 3,000; e.g. Visium data), uniformly distributed data points (*n* = 5,000; e.g. MERFISH data), and data points with a mixture of uniform and Gaussian distributions (*n* = 5,000) representing other complex data types. For gene expression of each data point, we randomly sampled from Gaussian distributions with varying levels of noise and separate distributions for active and inactive ligand and receptor genes. Using these benchmarks, we compared our model with 5 methods: NEST, COMMOT, Niches, K-nearest-neighbor-based distancing, rectified linear activation function for an alternative activation function, and a naive model (see Methods).

**Figure 3:**
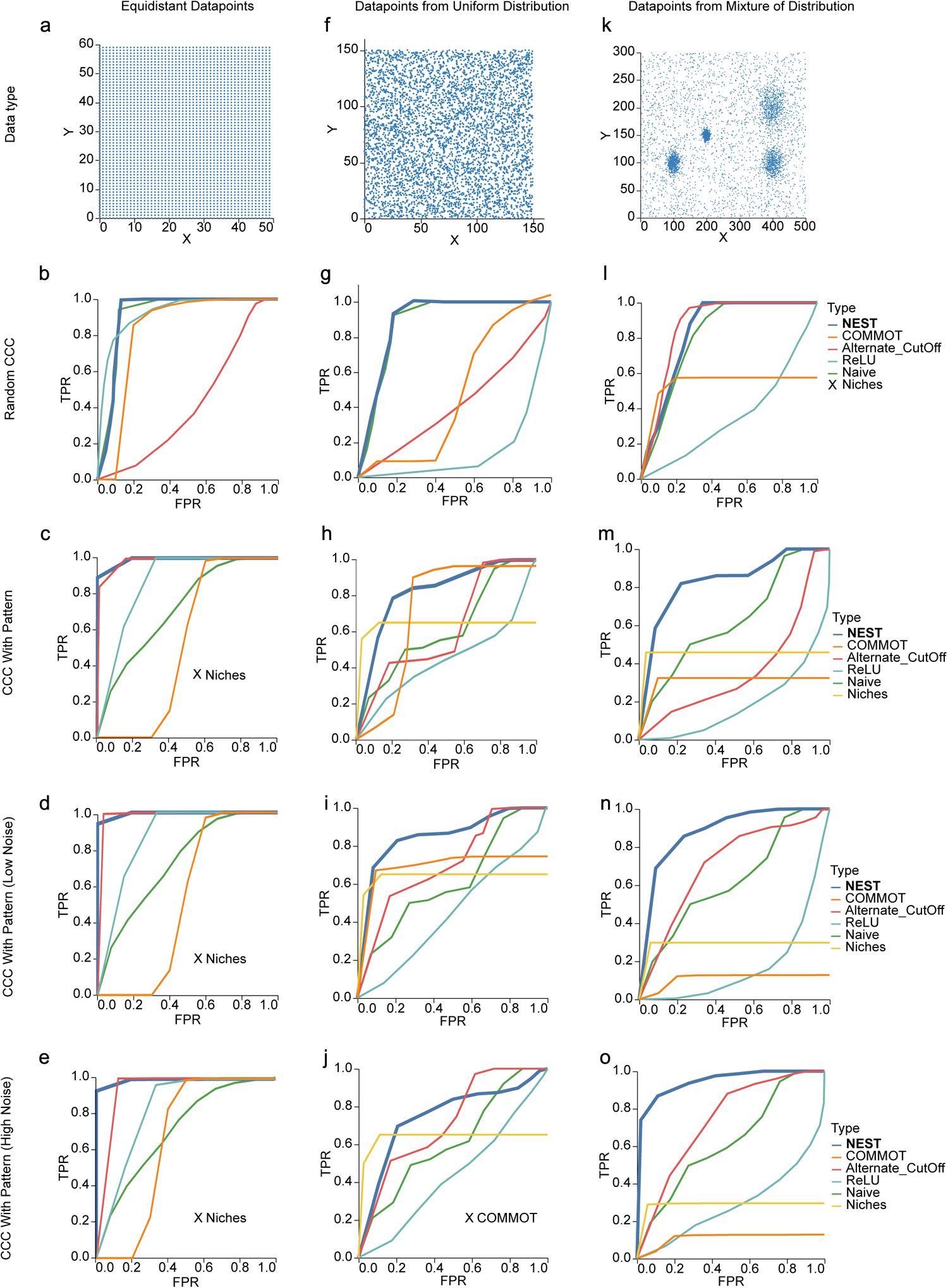
Quantitative performance analysis of NEST against other methods using synthetic benchmarks. Performance of NEST, Naive, ReLU (the NEST architecture with ReLU instead of the Tanh activation function), COMMOT, and Niches shown as ROC curves for equidistant (**a-e**), uniformly random (**f-j**), and mixture (**k-o**) of distributions. “X” in the legend indicates incomplete results due to error. **a** Data with equidistant data points representing Visium samples. **b-e** Performance of different models on equidistant data with randomly introduced synthetic CCC without forming any pattern (**b**), simulated patterns (**c**), patterns with low injected noise (**d**), and patterns with high injected noise (**e**). Niches did not run for **c-e** due to an error and thus marked as “X”. **f** Data with uniformly distributed data points representing MERFISH samples. **g-j** Performance measured as in **b-e** for the data in **f**. “X” marks COMMOT which did not detect any true positives CCC with high amount of noise, and marked with cross. **k** Data Points from a mixture of Gaussian and uniform distributions representing complex data format. **l-o** Performance measured as in **b-e** for the data in **k**.

Our benchmark for equidistant spot-based data demonstrated the advantage of NEST’s model over other methods, reaching the maximum true positive rate much faster and thus outperforming each algorithm (Figure 3a,b). Even so, this benchmark provides a basis for all methods, as they all tend to identify separate true ligand-receptor pairs from noise. However, since signals in biological samples tend to follow patterns rather than be defined by noise, we implemented a “pattern”-based benchmark. For example, if a type *s* signal is sent from a sender cell *i* to a receiver cell *j*, then *j* can send another type *t* signal in response as a sender cell to a receiver cell *k*. These “patterns” of communication enable multiple cells to execute a function collectively and serve as a valuable benchmark for detecting cell-cell communication^42^. When using a pattern-based benchmark, we observed a striking separation of methods, with NEST outperforming all other models across multiple levels of noise (Figure 3c-e). The naive model cannot detect such patterns by only prioritizing highly expressed genes of closely residing data points (Figure 3 and Supplementary Figure S1b-c). Niches and COMMOT mainly focus on differentially expressed and highly expressed ligand/receptor genes and do not recognize re-occurring patterns. Interestingly, the model with ReLU activation function does not perform well due to its unstable nature (Supplementary Figure S1d-e).

NEST’s performance is consistent in mostly outperforming other methods both in non-pattern- and pattern-based benchmarks in our uniform distribution benchmarks across all noise injections (Figure 3f-j and Supplementary Figure S1f). Surprisingly, this also holds true for the mixture of distributions as well, suggesting that NEST is robust and reliable for many different use cases and technologies and complements existing methods (Figure 3k-o).

### NEST identifies communication between single cells in mouse hypothalamus

Our synthetic benchmark demonstrated the capability of NEST to detect not only CCC in spot-based grids but also in distributions of single cells. To evaluate NEST’s performance on single-cell resolution spatial transcriptomic technologies, we applied NEST to MERFISH slides from the hypothalamus preoptic region of female parent (*n* = 5,533) and female virgin (*n* = 5,606) mice to investigate differences in CCC^34^. NEST revealed that female parent and virgin tissues varied in spatial distributions of strong communication (Figure 4a-b). In concordance with previous findings, NEST identified the neuropeptide galanin as *GALR* in both parent and virgin mice, along with the brain-derived neurotrophic factor *BDNF* through *BDNF*-*ESR1* for warm temperature sensitiveness^34,43^ (Figure 4c-d). In contrast, NEST mainly detected oxytocin and receptor genes *OXT* and *OXTR* in the female parent mouse only, which are part of the core parenting signals^44,45^ (Figure 4c).

**Figure 4:**
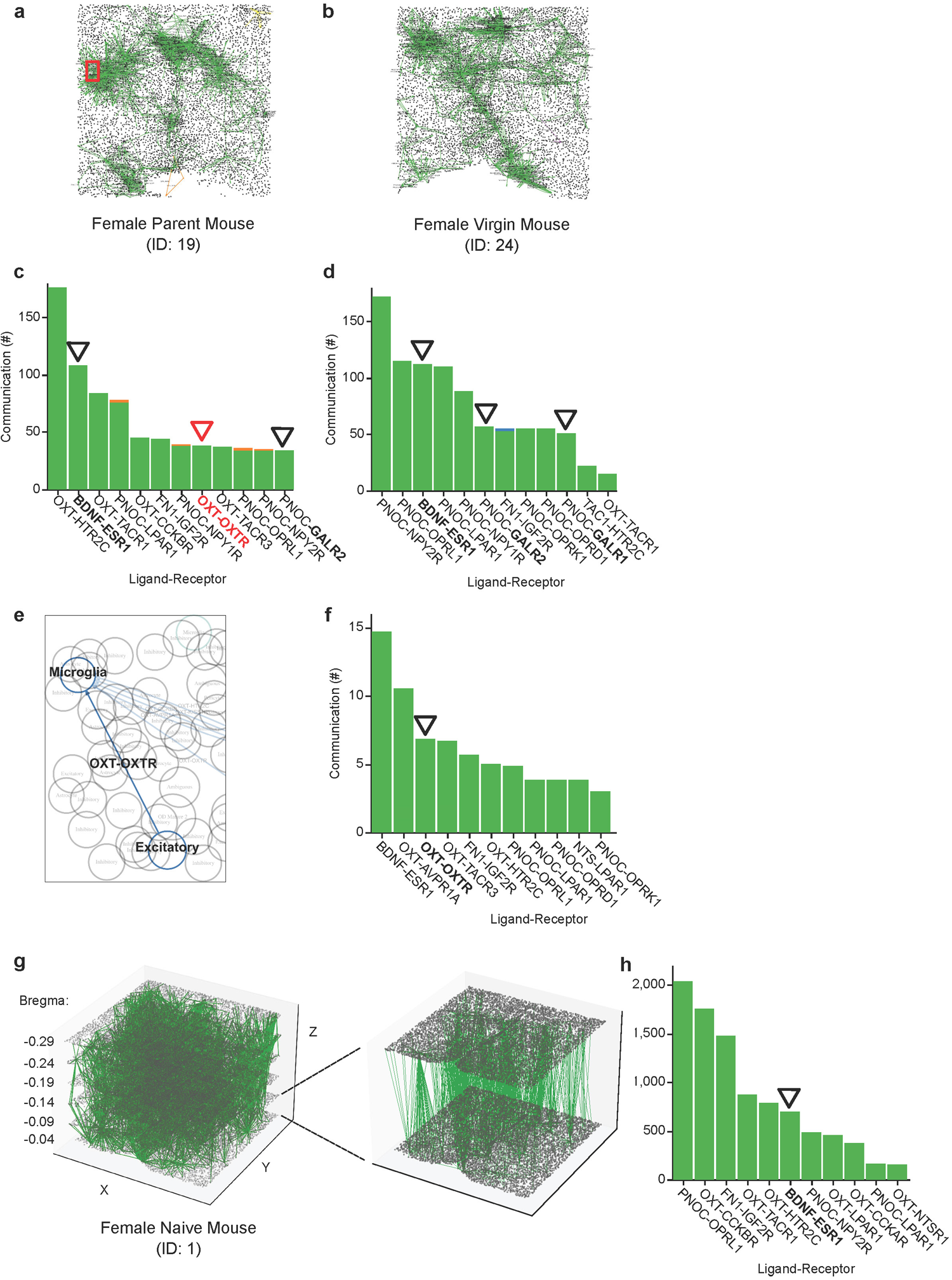
NEST identifies communication involved in mouse parental behavior from the hypothalamus preoptic region measured with MERFISH. **a-b** NEST detected communication (green) in tissue from female parent mouse (**a**) and female virgin mouse (**b**). **c-d** Corresponding histograms for the parent mouse (**c**) and virgin mouse (**d**) tissues showing abundance of signals of the top 20% strongest ligand-receptor pairs. NEST identified the neuropeptide galanin as *GALR* in both parent and virgin mice, along with the brain-derived neurotrophic factor *BDNF* through *BDNF*-*ESR1* for warm temperature sensitiveness (marked with black triangle). In contrast, the parenting signal *OXT* -*OXTR* is exclusively found in the female parent mouse (red triangle). **e** Cell-type specific communication zoomed in from **a** (red rectangle). **f** The corresponding NEST-generated histograms from **e** showing CCC between microglia and excitatory neurons. As in **c**, *OXT* -*OXTR* is detected (black triangle) as one of the strongest communications that contributes to emotional bonding within the female parent mouse. **g** 3D MERFISH sample from a female naive mouse (ID: 1) with NEST-detected communication (left), with a zoom-in of two layers for clearer visualization (right). **h** Histogram of communication found in **g**.

Importantly, we found that NEST could detect communication between two individual cells: a neuron and a microglial cell (Figure 4e,f). Using the single-cell MERFISH female parent mouse sample with previously annotated cell types, we filtered ligand-receptor pairs identified by NEST such that the sender and receiver cells were classified as neuron or microglia only. Upon inspection, we observed a strikingly high-resolution image where we could observe a predicted single excitatory neuron sending an *OXT* signal to a *OXTR* receptor on a receiving microglia (Figure 4e). This ligand-receptor pair establishes communication that contributes to emotional bonding within the female parent mouse^46^. This communication was well-represented across all neuron-microglia communication (Figure 4f). Together, this analysis suggests that NEST can detect more precise cell signaling at single-cell resolution rather than solely between pseudobulk cell types.

Spatial transcriptomic technologies such as MERFISH also may take consecutive slices to infer three-dimensional (3D) cell organization (Figure 4g). We sought to extend our model for 3D data points by combining *n* = 38,372 cells across 6 such consecutive slides along the bregma axis. As NEST uses a graph structure that is not limited to two dimensions, we extended our edges to incorporate 3D input where the physical distance matrix records pairwise distances of 3D coordinates. When applying NEST on a 3D female naive mouse sample, NEST detected general communication in the mouse brain while suppressing parental signals, likely because this mouse was not exposed to pups^34^ (Figure 4g,h). Applying NEST to 2D and 3D samples revealed the unique flexibility of NEST across technologies and dimensions.

### NEST detects aggressive cancer CCC in lung adenocarcinoma

The tumor microenvironment is a complex and heterogeneous collection of different cell types and signals, where CCC contributes to disease progression. To identify specific regions of tumor tissue associated with cancer-promoting communication, we applied NEST to a Visium sample of lung adenocarcinoma (LUAD) containing 4,095 spots^33^ (Figure 5a). Within the most probable ligand-receptor pairs, NEST detected transforming growth factor *β* genes *TGFB1* and *TGFB2*, important in metastasis in LUAD^47^, as concentrated near the top out of over 12,605 pairs in the database based on attention scores (Figure 5b). However, NEST found apolipoprotein E (*APOE*)-based communication including *APOE*-*SDC1* as the most strongly occurring CCC. APOE promotes lung adenocarcinoma proliferation, migration, and is considered as a potential survival marker in lung cancer^48^. To confirm the presence of *APOE*-*SDC1* within the tumor region, we observed alignment between the expression of each gene on the tissue with the location of the CCC (Figure 5c-e).

**Figure 5:**
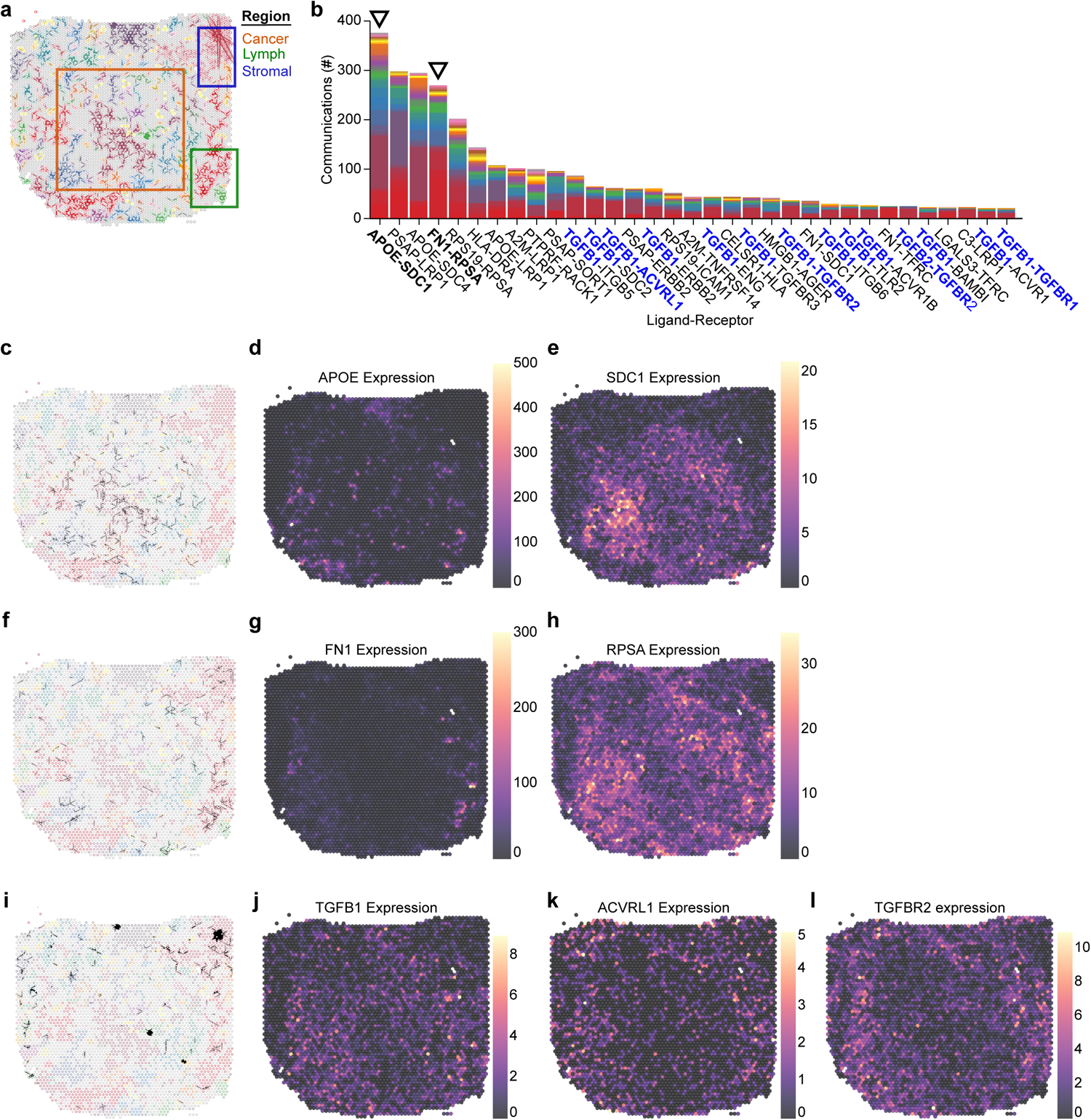
NEST detects localized signaling in tumor and stromal environments in LUAD. **a** NEST-generated communication graph showing regions with strong CCC colored by component. Gray indicates regions with no or weak CCC. Top right red component has thinner arrow widths to accommodate high communication frequency between a few spots. The boxes outline three regions: cancer (orange), lymph (green), and stromal (blue) based on prior histological annotations (generalized from Zhu et al.^33^). **b** Histogram collecting ligand-pair receptor abundance (*y*-axis) from **a**, colored by connected component. The *APOE*-*SDC1* and *FN1*-*RPSA* signals (marked with triangles) are exclusively detected by NEST, which are associated with LUAD proliferation and lymph node metastasis. NEST also detects many *TGFB* (blue text) signals that causes metastasis in LUAD. **c-l** Location of specific tumor and stroma signals found in **b**. **c** Communication from **a** filtered for *APOE*-*SDC1* signals. This is the most abundant signal and is mainly found in cancer-annotated regions. **d-e** Gene expression of *APOE* (**d**) and *SDC1* (**e**) on **a**, mainly in cancer regions. **f** Communication from **a** filtered for *FN1*-*RPSA* signals. This signal maps to the lymph nodes. **g-h** Gene expression of *FN1* (**g**) and *RPSA* (**h**) on **a**. NEST detects this ligand-receptor pair mainly in the bottom right lymph node region where both genes are highly expressed. **i** Communication from **a** filtered for *TGFB1* signals. **j-l** Gene expression of the *TGFB1* ligand (**j**), *ACVRL1* (**k**), and *TGFBR2* (**l**) on **a**. These signals are more common within the tumor microenvironment than the tumor region.

In addition to tumor-localized *APOE-SDC1*, NEST identified other strong communication in different locations of the tissue. Specifically, NEST assigned *FN1*-*RPSA* to the lymph node (Figure 5f-h). As *FN1* is associated with lymph node metastasis as well as *APOE* in patients with LUAD, NEST may have identified potential disease progression^49,50^. Separate from the lymph node, NEST identified *TGFB* signaling within the surrounding tumor microenvironment^51,52^ (Figure 5i-l). Based on these observations, NEST is able to deconvolve complex tumor microenvironments to place signals in precise regions of tissue.

### NEST identifies consistent ligand-receptor pairs between pancreatic cancer tissues

To see if NEST would generalize to other cancer types with heterogeneous regions, we applied NEST to pancreatic ductal adenocarcinoma (PDAC) tissues. PDAC is widely recognized as a highly aggressive disease, yet treatment responses can vary widely among patients. There is immense transcriptional diversity defining discrete “Classical” and “Basal” subtypes of PDAC that is crucial in explaining treatment heterogeneity. Basal tumors exhibit characteristics reminiscent of basal or squamous epithelium, leading to heightened chemoresistance and poorer patient prognosis. Conversely, the Classical tumors demonstrate transcription factor expression associated with pancreas development, rendering them more responsive to chemotherapy and yielding improved clinical outcomes^34,53–55^. As the PDAC tumor microenvironment is a heterogeneous and dense collection of tumor, stromal, and immune cells, clear subtype assignment within a tissue is challenging. Different types of stromal areas with high (activated) or low (deserted) immune activity contribute to divergent regions within tumor tissue^56^. To date, the relationship between divergent regions, transcriptomic subtypes, and cell states of PDAC is unclear. To resolve specific cell-cell interactions in this complex disease, we applied NEST which considers high-resolution tumor and stromal proximity and does not rely solely on highly expressed genes. We tested whether NEST could find CCC separated by the spatial location of PDAC transcriptomic subtypes.

We applied NEST to Visium data collected from two cases that showed morphological heterogeneity across tissue regions (Figure 6). Transcriptomic subtypes are known to correlate with tumor morphology. Classical tumors are well differentiated and have a gland-forming morphology, while Basal tumors are moderately to poorly differentiated with non-gland forming morphology^57,58^. Both cases were resectable, stage IIb PDAC tumor samples (PDAC_64630 and PDAC_140694). Sample PDAC_64630 (*n* = 1,406 spots) presented several regions of morphologically and transcriptionally distinct tumor subtypes split by stroma^57^ (Figure 6a-b, see Methods).

**Figure 6:**
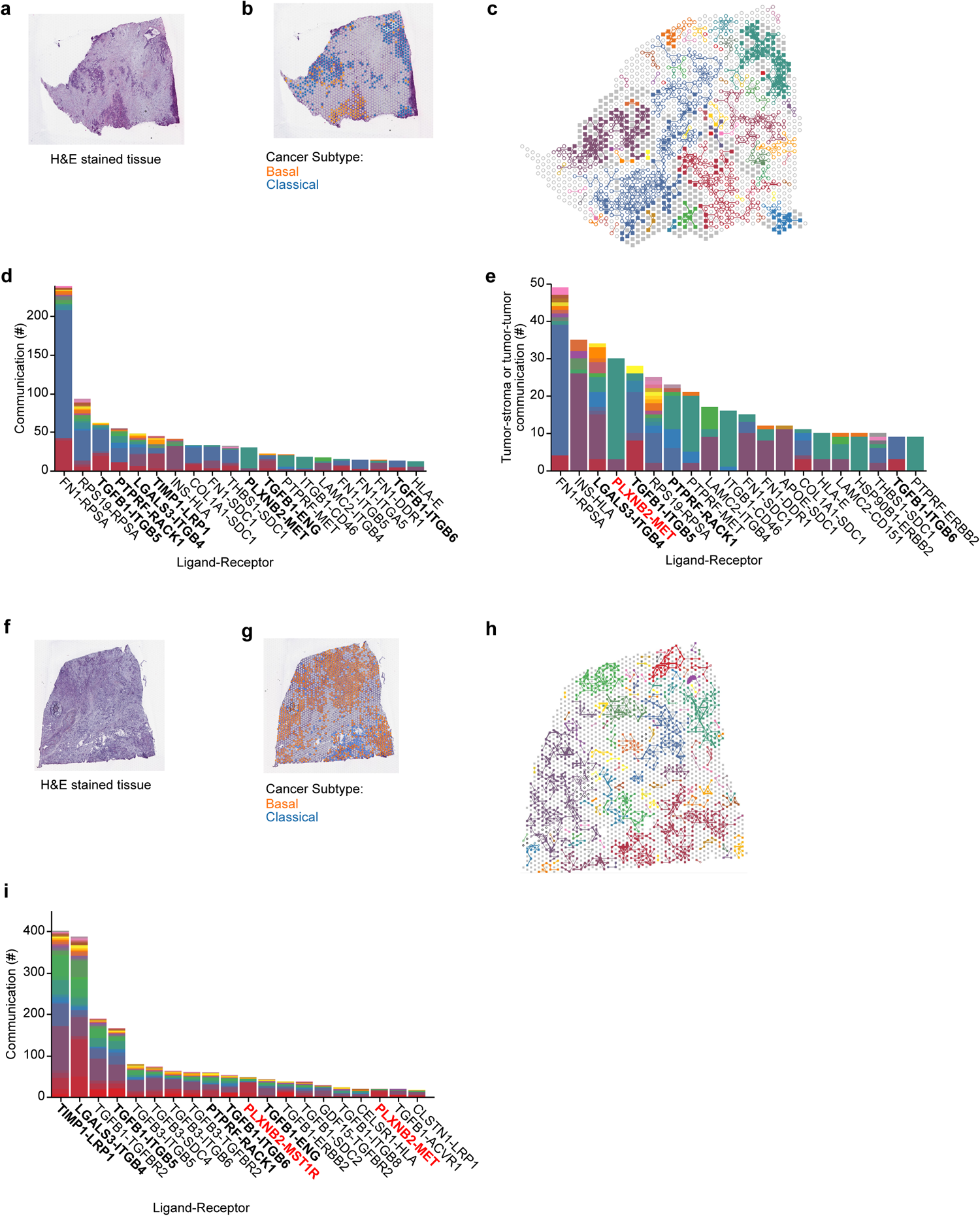
NEST reveals subtype-specific patterns of communication in PDAC tissue. **a-b** Tissue from PDAC_64630 measured by Visium with an hematoxylin and eosin stain (H&E; **a**) or colored by PDAC subtype (**b**) determined by gene signatures. **c** NEST-detected communicating regions from **a**. NEST-produced output graphs show tumor (filled square) and stroma (open circle) regions. Colors correspond to different connected components. **d-e** Histogram of the strongest CCC counts (*y*-axis) determined by NEST across the whole tissue (**d**) or tumor-communicating regions (**e**). Colors correspond to each connected component in **c**. **f-g** Tumor tissue from patient PDAC_140694 measured by Visium with H&E (**f**) or colored by PDAC subtype (**g**) determined by gene signatures. **h-i** NEST-detected strongly communicating regions from **f** (**h**) with corresponding histogram of CCC counts (**i**) colored by each connected component in **h**. Ligand-receptor pairs highlighted with bold text in **d** and **i** represent a set of communications that is detected in both samples with NEST. The *PLXNB2*-*MET/MST1R* highlighted with red text in **e** and **i** represents a communication that is associated with the Classical subtype in both samples by NEST.

We first assessed if NEST could identify PDAC-relevant ligand-receptor pairs across the whole tissue. NEST reported 411 ligand-receptor pairs out of 12,605 total pairs in the top 20% strongest signals, with *FN1*-*RPSA* as the most abundant with an occurrence of 239 instances, which was mainly found in the stroma region (Figure 6c-e and Supplementary Figure S2a). Fibronectin (*FN1*) is considered one of the main extracellular matrix constituents of pancreatic tumor stroma. High stromal *FN1* expression associates with more aggressive tumors in patients with resected PDAC^59^. Likewise, the cell membrane receptor Ribosomal Protein SA (*RPSA*) regulates pancreatic cancer cell migration^60^. We observed additional canonical signals in pancreatic cancer such as *TGFB* signaling, which promotes fibrosis and immune evasion^61^, and Protein Tyrosine Phosphatase Receptor Type F (*PT-PRF*), which contributes to cancer-cell invasiveness and is under active consideration as a therapeutic target for pancreatic cancer^62^.

To determine whether NEST could identify consistent tumor-associated CCC across multiple tissues, we applied our model to PDAC_140694 (*n* = 2,298 spots) derived from a different patient with similar PDAC subtypes to PDAC_64630 (Figure 6f-i). PDAC_-140694 contained mostly tumor cells, with less stroma compared to PDAC_64630. To directly compare communication occurring within each sample, we filtered NEST-identified signals in PDAC_64630 to those between tumor spots only or tumor and stromal spots (Figure 6e,h-i). We found overlapping PDAC-associated CCC between both patients in the top 20 strongest signals, including *LGALS3*-*ITGB4*, *PLXNB2*-*MET* /*MST1R*, *PTPRF*-*RACK1*, *TGFB1*-*ITGB5*, and *TIMP1*-*LRP1*^63^ (Figure 6e,i). The high concordance of top signals suggests NEST can detect similar communication in similar contexts.

### NEST reveals PDAC subtype-region-specific communication

After identifying tumor-wide CCC associated with PDAC, we tested if NEST could resolve CCC within specific tissue regions. We annotated tumor regions according to Classical and Basal transcriptomic PDAC subtypes^64^. Using NEST, we visually detected region-specific CCC *PLXNB2*-*MET* /*MST1R* primarily in the Classical region across both samples (Figure 6c,e,h-i and Supplementary Figure S2b-g). Furthermore, we observed a correlation of CCC frequency with CCC type (Chi-square test of dependency: *p* = 6.14 × 10^−32^) and significantly higher occurrence of *PLXNB2*-*MET* over all other communications in the Classical region (Hypergeometric test of over-representation: *p* = 1.41 × 10^−22^; Supplementary Note S1). *PLXNB2* codes for a plexin protein, part of a family of transmembrane receptors, initially recognized for their role in axon guidance. Plexins are known for their key role in tumor CCC, tumor growth, migration, and metastasis. Semaphorins are the main ligands of the Plexin receptors; however, some plexins can also form complexes with other tyrosine-kinase receptors, such as the hepatocyte growth factor receptor (HGFR) encoded by *MET* ^65^ and RON encoded by *MST1R*^66^. NEST reported a *PLXNB2*-*MET* /*MSTR1* CCC pattern that is favored in Classical over Basal areas of the tumor. To further explore differences in the *PLXNB2*-*MET* axis between Classical and Basal tumor cells, we used RNA sequencing data from a library of 10 PDAC patient-derived organoid models (Figure 7a-d). Organoid gene expression confirmed that Classical tumors exhibit significantly higher *MET* expression than Basal tumors (Fisher-Pitman permutation test: *p* = 3.18 × 10^−2^; Figure 7a-b, Supplementary Note S2). While the role of plexins are described in other solid tumors including PDAC, most of the studies explored semaphorins as their predominant ligands^67^. Here, NEST suggests an underexplored subtype-specific CCC pattern.

**Figure 7:**
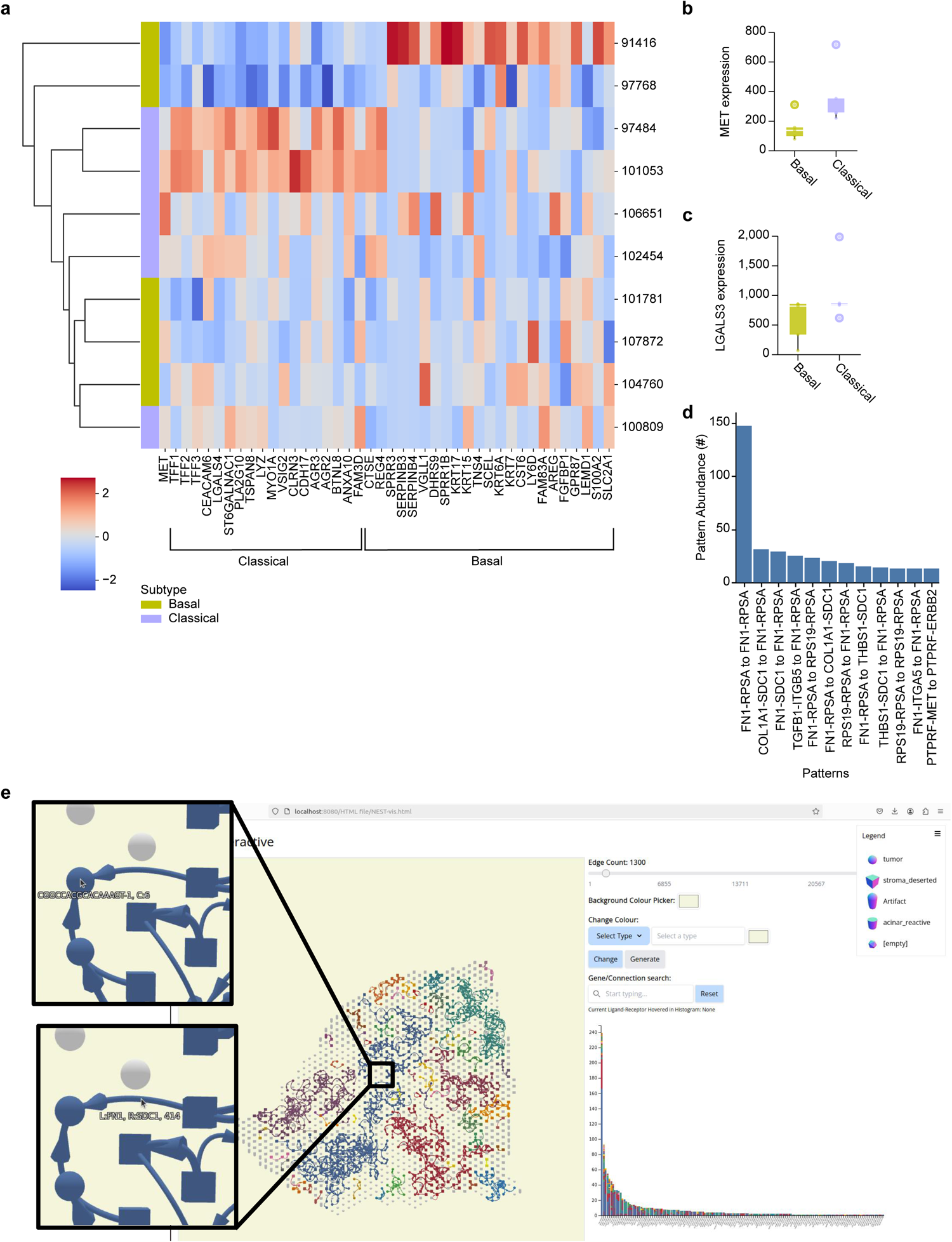
**a** Heatmap displaying gene expression of *MET*, Classical-associated genes, and Basal-associated genes^64^ in PDAC patient-derived organoid models (*n* = 10) derived from bulk RNA-sequencing. **b-c** Box and whisker plots comparing gene expression derived from bulk RNA-sequencing within Basal (*n* = 5) versus Classical (*n* = 5) patient-derived PDAC organoids classified with an established subtyping scheme^64^. Center line: median; box: interquartile range; whiskers: 1.5 ×interquartile range; points: outliers. **b** *MET* expression is significantly higher in the Classical subtype (Fisher-Pitman permutation test: *p* = 3.18 × 10^−2^). **c** *LGALS3* expression is found in both Basal and Classical subtype (Fisher-Pitman permutation test: *p* = 0.175). **d** Frequency of two-hop CCC patterns (ligand-receptor-ligand-receptor) in sample PDAC_64630. **e** Overview of NEST Interactive. The NEST Interactive display shows a fully interactive network of vertices (cells or spots) connected by ligand-receptor pairs (left). The display features a user interface (right) including options for filtering genes, selecting thresholds for attention scores of communication, as well as a histogram based on the location of each ligand-receptor pair. Highlighted in zoomed insets is a stroma-deserted vertex sending ligand *FN1* (bottom) to a tumor which is receiving the signal through its *SDC1* receptor (top).

In contrast to *PLXNB2*-*MET*, NEST detected *LGALS3*-*ITGB4* in Basal and Classical mixed regions. Galectin-3 (*LGALS3*) mediates tumor-stroma interactions by activating pancreatic stellate cells^68^. We observed equally high expression of LGALS3 in both Classical and Basal organoids (Fisher-Pitman permutation test: *p* = 0.175; Figure 7c and Supplementary Note S2). To determine the potential impact of these signals, we explored the association between these genes and PDAC using The Cancer Genome Atlas^69^. All NEST-identified genes were classified as “unfavorable” in the context of PDAC, and associated with survival (Log-rank test: *p* = 0.014,0 for *FN1*, *p* = 8.50 × 10^−3^ for *PLXNB2*, *p* = 1.21 × 10^−7^ for *MET*, *p* = 6.39 × 10^−4^ for *ITGB4*, and *p* = 1.40 × 10^−4^ for *ITGB5*). Furthermore, *MET*, *ITGB4*, and *ITGB5* achieved high antibody staining results for PDAC and were considered prognostic by the Human Pathology Atlas^70^ which highlighted these genes as potential targets for treatment. Importantly, NEST’s top identified ligand corroborates previous findings illustrating the critical role of *FN1* as a pivotal signaling gene against pancreatic cancer based on survival and gene expression analyses^71^. Together, these findings suggest that different subtypes of PDAC use distinct tumor-promoting CCC that impact patient outcome.

NEST’s unique capability extends single ligand-receptor pairs to patterns of communication as a relay network. We next sought to characterize differences between our previously identified ligand-receptor pairs with patterns within pancreatic tumor tissue. Although NEST detects any type of patterns, for simplicity we counted the frequency of two-hop CCC between cells (Figure 7d). Specifically, a ligand-receptor pair *s* from an *i* sender to a *j* receiver, and ligand-receptor pair *t* from *j* sender to *k* receiver. NEST detected *FN1-RPSA to FN1-RPSA*, *COL1A1-SDC1 to FN1-RPSA*, *TGFB1-ITGB5 to FN1-RPSA*, and *PLXNB2-MET to PTPRF-MET* among the most frequently occurring pattern of this type (Figure 7d). These signals promote cell adhesion (*FN1* and *TGFB1*^63^), migration (*RPSA*^60^), metastasis (*FN1*^71^), collagen proteins (*COL1A*^72^), and inflammation (*SDC1*^73^). Unlike single ligand-receptor pair analyses, this pattern identifies many inflammatory pathways, suggesting a cascading pattern that could not have been detected without NEST.

### NEST Interactive provides a web-based visualization tool for exploring cell-cell communication

To help visualize cell-cell communication on tissues, we developed NEST Interactive as a web-based data visualization tool (Figure 7e and Supplementary Figures S7 and S8). NEST Interactive features a 3D, responsive graph illustrating cells or spots as vertices and ligand-receptor pairs as directed edges. The user is able to specify the number of strongest ligand-receptor pairs which updates connected components and colors on-the-fly. NEST Interactive also displays a corresponding histogram listing each unique ligand-receptor pair stacked by connected components showing their specific region of tissue. The user can visualize a particular gene or ligand-receptor pair on both the 3D graph and the histogram using a fuzzy search feature. NEST Interactive is designed with responsiveness in mind for both mobile and PC. NEST Interactive is available at https://github.com/schwartzlab-methods/NEST-interactive.

## Discussion

Detecting communication through ligand-receptor interaction is necessary to decipher cellular activity in tissue. Existing scRNA-seq-based computational methods for identifying CCC in tissue samples often produce an extensive number of false positives by lacking cell-cell proximity information. Recent spatial transcriptomics-based tools either quantify CCC at cell-population resolution, missing critical rare communication events, or do not consider patterns of ligand-receptor usage. We overcome these challenges by introducing NEST, which integrates ligand-receptor information with cell location through a graph attention network at single-cell resolution. We quantitatively evaluated NEST and found our model to have superior performance against other available methodologies using new benchmarks of 12 different arrangements of synthetic data representing different technologies and species. To further validate and identify new aspects of disease, we applied our model to biological samples at different resolutions and dimensions under healthy and diseased conditions. NEST consistently captured known CCC on all samples. We pinpointed distinct CCC between pancreatic ductal adenocarcinoma subtypes that were consistent between patients and associated with overall survival from independent cohorts from The Cancer Genome Atlas and Human Protein Atlas.

Existing spatial transcriptomic methods for detecting CCC, such as COMMOT and Niches, focus on high co-expressions of ligand-receptor pairs and do not attempt to recognize patterns of activity. Patterns may also correlate with tissue regions even when lowly expressed. Using a pattern recognition algorithm may contribute to NEST’s advantage over other methods when identifying T-cell homing signals in precise locations of T-cell zones in human lymph nodes. Moreover, NEST uses all genes to identify CCC, orders communication based on learned importance, and spatially pinpoints their location —not filtering out low variance ligand-receptor pairs which would prevent the method from detecting genes that are well-characterized and a part of informative modules, but with low variability^74^. NEST identified expected signals that were not among the most highly expressed, indicating the importance of integrating spatial and molecular information. These unique capabilities of NEST help associate CCC with the target disease and their subtypes.

Recent methods alternatively use scRNAseq data for CCC detection before mapping ligand-receptor pairs to spatial data^15,16,33^. However, such tools only resolve CCC between adjacent cells or spots and do not discriminate between distant ligand-receptor mechanisms such as paracrine, which constitute the majority of ligand-receptor databases. In contrast, NEST incorporates three major types of communication: autocrine (self-communication), juxtacrine (communication between adjacent cells), and paracrine (communication between nearby, non-adjacent cells). Furthermore, existing machine-learning-based tools such as GraphComm^16^ use supervised learning, which is difficult to train due to the unavailability of labeled data. To address this issue, NEST applies contrastive learning, an unsupervised training to overcome the ground truth data scarcity problem. This powerful and generalizable architecture renders NEST applicable to a wide range of data based on resolution, two or three dimensions, and healthy or diseased conditions.

To enable this flexibility, NEST only requires a ligand-receptor database for *a priori* knowledge. We acknowledge that NEST may produce some false positive results in noisy conditions due to other factors that impact CCC in a tissue. For example, intracellular signals both trigger ligand production and determine cell-surface receptor expression. Gene regulatory networks in a cell work to synchronize cell actions and may influence CCC. Future studies may extend NEST to model ligand-receptor-target signaling networks using an intracellular network and gene regulatory networks^11,75^. To assist with intracellular measurement, subcellular information from MERFISH and Xenium experiments may contribute to the model. With a mix of transcripts of genes involved in intra- and intercellular communication, a restructured model could attain higher CCC detection accuracy.

With the advancement of spatial-omics technologies, future models may incorporate other data modalities such as proteins using currently available Xenium data, Visium HD, or chromatin accessibility from emerging assays^76^ to improve CCC detection. NEST’s underlying model is flexible with the expectation of integrating additional data types. We anticipate methods like NEST that take full advantage of the spatial proximity of cells will reveal new avenues of determining cellular neighborhoods and their contribution to health and disease.

## Methods

### NEST architecture

NEST is an end-to-end solution for processing data directly from a spatial transcriptomic data structure from programs such as Space Ranger, detecting strong signals and patterns of communication within specific regions of tissue, and displaying CCC through an accessible visualization. There are four main steps in the NEST workflow: Data Preprocessing Step, Input Graph Generation Step, Communication Prediction Step, and Output Graph Generation Step (Figure 1).

### Data Preprocessing Step

NEST takes four inputs: a spatial transcriptomic data, a ligand-receptor database, a threshold percentile *t_g_* (e.g. 80^th^ or 98^th^ percentile) to select highly expressed genes, and a threshold distance *t_n_* as a neighborhood cut-off distance (Figure 1a-c). The default database provided by our model is a combination of the CellChat and NicheNET databases, totaling 12,605 ligand-receptor pairs. For *N* spots or cells (here called vertices) and *M* genes in a spatial transcriptomic data set, NEST generates a gene expression matrix **A** ∈ ℝ*^N^*^×^*^M^*. NEST calculates the Euclidean distance between each pair of vertices to generate a physical distance matrix of dimension **D** ∈ ℝ*^N^*^×^*^N^*. NEST uses quantile normalization^77,78^ on the gene-expression matrix to standardize gene distributions across cells or spots to enable direct comparisons. For each cell or spot in the gene expression matrix, NEST considers genes having expression over *t_g_* percentile (default 98) as active.

### Input Graph Generation Step

After preprocessing, NEST generates an input graph *G* = (*V*, *E*), where *V* (|*V* | = *N*) represents the set of vertices (cells or spots) and *E* (where |*E*| is typically over 1 × 10^6^) represent the set of neighborhood relations among the vertices in *G* (Figure 1d). We add a neighborhood relation between a vertex *i* and *j* if the distance between *i* and *j* is less than or equal to *t_n_*. For each ligand *l* and paired receptor *r* from the ligand-receptor database, if **A***_i_*,*_l_* and **A***_j_*,*_r_* are active, NEST will insert a directed edge from *i* to *j*. NEST allows for multiple edges to represent multiple ligand-receptor pairs between two vertices. Importantly, an edge between a pair of vertices does not necessarily mean that a communication is happening along that edge because CCC communication is highly context dependant^13^ and effected by various epigenetic factors^37^. An edge is just neighborhood relation representation between a pair of vertices which NEST will decide is a probable CCC.

We next pass *G* to the deep learning module “Communication Prediction Step” through two input feature matrices: a vertex feature matrix 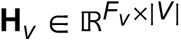 and an edge feature matrix 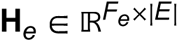. Each column in **H***_v_* is a vertex input feature vector (e.g. 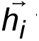 for vertex *i*), which represent each cell or spot in the dataset. NEST uses a one hot vector to uniquely present each vertex, so *F_v_* = |*V* | (Figure 1d). Similarly, each column in the edge feature matrix, **H***_e_*, is an edge feature vector representing an edge (neighborhood relation) in *G*. The edge feature vector has dimension *F_e_* = 3 since it has three attributes (Figure 1d): physical distance between vertices (e.g. *d_i_*,*_j_* from the physical distance matrix **D** ∈ ℝ*^N^*^×^*^N^*), ligand receptor co-expression score for the corresponding edge (e.g. *L_e_* × *R_e_* from Figure 1c), and the identifier of that ligand-receptor pair from the input database (Figure 1b). We pass these two input feature matrices to the next step, the “Communication Prediction Step”.

### Communication Prediction Step: Overview

The NEST architecture builds on two main deep learning concepts: graph attention networks^36^ (GAT) as encoders and deep graph infomax^29^ (DGI) to train encoders through contrastive learning (Figure 1e). Although based on GAT models that are traditionally used with a training set, there is no ground truth for CCC detection, so NEST instead uses DGI for training. We provide implementation functions for integrating GAT into the DGI model in our Github repository located at https://github.com/schwartzlab-methods/NEST/blob/main/CCC_gat.py.

### Communication Prediction Step: Graph attention network as an encoder

The GAT generates a vertex embedding which encodes information about a vertex *i* in *G* along with its neighborhood information, here meaning which vertices can *i* communicate with and through which ligand-receptor pairs. The attention module in the GAT assigns “attention scores” to each edge based on how necessary and sufficient those edges are to capture hidden patterns that together reconstruct the input sample.

Let input vertex feature vectors for vertices *i* and *j* be 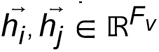, input edge feature vectors from *j* to *i* be 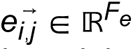, and the dimensions of vertex and edge embeddings be *F*^′^. Then the learnable weight matrix for the linear transformation of vertex features is 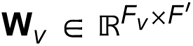, while the equivalent matrix for edge features is 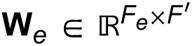. Then the attention score for the edge from *j* to *i* is

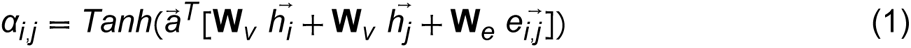

This score indicates the importance of vertex *j*’s features to vertex *i*. Here, the attention 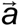 is a learnable parameter where 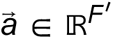. Here we use Tanh, as we found increased performance using Tanh non-linearity instead of PReLU (and ReLU), the latter of which was too unstable (Figure 3 and Supplementary Figure S1d-e). After learning the attention scores, we apply a Softmax normalization over all incoming edges to vertex *i* from its neighbors *N_i_* using

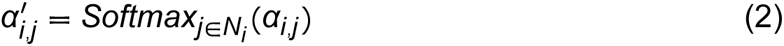

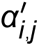 ranges between 0 and 1 in an effort to scale attention scores. We use Softmax normalization for the message propagating principle. Using the normalized attention scores, we obtain a vertex embedding for *i* with

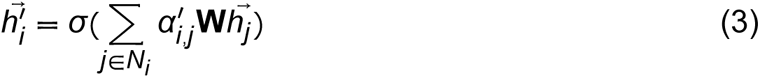

Here, the GAT generates a vertex embedding matrix 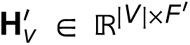. However, for communication prediction, we use the attention scores rather than the vertex embedding to prioritize edges in set *E* based on global context. To detect which regions are more active than others in the input sample, we use unnormalized attention scores from Equation (1), as these scores are globally comparable across the tissue (Supplementary Figure S3a). As such, we use the scores obtained by Equation (1) directly as a representative of cell-cell communication probability. We can scale these scores between 0 to 1 over all the edges in *E* such that scores closer to 1 present higher probability of communication.

NEST generally assigns higher attention scores to input edges with high ligand-receptor coexpression scores (Supplementary Figure S3b-m). Importantly, the conventional way of using normalized attention scores cannot achieve this goal (Supplementary Figure S3a), so NEST uses unnormalized attention scores assigned by the GAT.

### Communication Prediction Step: DGI for encoder training

We apply the contrastive learning model DGI^29^ to train the GAT in an unsupervised approach. DGI takes the input graph *G* = (*V*, *E*) and applies random permutation, shuffling edges to form a corrupted graph *G_C_* = (*V*, *E*^′^), where *E*^′^ is the set of corrupted edges (Supplementary Figure S1a). We store the original input graph as *G_T_*. This contrastive learning approach has two branches to handle each version of the input graph: the corrupted branch and the original branch.

Both branches use the same GAT encoder, with shared learnable parameters or weight matrices, to generate a vertex embedding matrix 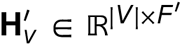. The vertex embedding generated from *G_T_* through original branch is summed to obtain the “summary vector” 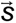. This summary vector captures global information content of the entire graph. We use a discriminator function to measure distance between 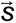 from the corrupted graph embedding (negative sample) and the true graph embedding (positive sample). NEST maximizes the mutual information between the summary vector and vertex embedding from the true graph by optimizing the Jensen-Shannon divergence between the negative and positive graphs. This divergence distance is related to the generative adversarial network distance^29^. Through many iterations (approximately 60,000 in our testing), NEST eventually converges to a minimal loss, and we save that model state.

### Output Graph Generation Step: Overview

NEST uses stochastic optimization Adam^79^ which may introduce small variations in the output of multiple runs. As an optional step to increase the accuracy and stability of communication detection, we run each experiment multiple times (default 5) with different seeds and combine the results from each run. Then, we apply postprocessing on the aggregated result to obtain the final output graph (Figure 1f).

### Output Graph Generation Step: Ensemble of multiple runs

We obtain the ranks of edges based on the attention scores assigned by each encoder layer for each run. Using the rank product^80^, we sort by the aggregated rank for each layer. We then merge results for both attention layers, as existing metapath work on graph neural networks suggests important characteristics are present in each layer^81^. This step also accepts a top percentage of communications *t_e_* as input from the user. By default, we select the top *t_e_* = 20% as the most reliable signals for the analyses presented here. We select this threshold on both layers independently.

### Output Graph Generation Step: Postprocessing for better visualization

This step postprocesses the list of strong CCC for better visualization and downstream analysis. We apply a connected component finding algorithm^38^ on the strongly communicating *t_e_* edges to generate subgraph components. In this way, we observe subgraphs where all vertices are strongly talking to at least one other vertex in the community, suggesting a set of vertices localized to specific regions. We provide several visualization outputs to best quantify NEST’s predictions, including a flexible dot file for plotting the graph, histograms delineating CCC within each region of tissue, and more (Figure 1g-h). We provide functions for this step at https://github.com/schwartzlab-methods/NEST/blob/main/output_visualization_NEST.py.

### Choosing model hyperparameters

NEST contains a complex model with rigorously tested hyperparameters. We held a comparative analysis of the loss versus iteration of NEST for different hidden layer sizes ranging from 64 to 2048 using PDAC_64630 sample (Supplementary Figure S4). 64 neurons had the longest convergence time and the component plot of the top 20% strongest ligand-receptor pairs did not align with cell-type annotation (Supplementary Figure S4b-c and Figure 6b). Both 512 and 2048 neurons provided similar component plots and CCC proportions, converging with a similar number of iterations (Supplementary Figure S4a,d-g). Therefore, we chose 512 as the hidden layer dimension to reduce GPU memory consumption while maintaining a good detection rate.

We also select the minimum number of cells required to keep a gene `min_cell` (by default 1), threshold percentile expression for gene selection *t_g_* (by default 98%), learning rate *l_r_* (by default 1 × 10^−5^), top-most signals to output *t_e_* (by default 20%), top count of edges `count`_edge_ for a more restrictive version of CCC selection during postprocessing (by default 1,500). A complete list of hyperparameters used for all the data sets presented in this paper are provided in Table 1. Interested users can change other hyperparameters related to GAT (e.g. number of attention heads) or DGI (e.g. discriminative function) according to their needs.

**Table 1:**
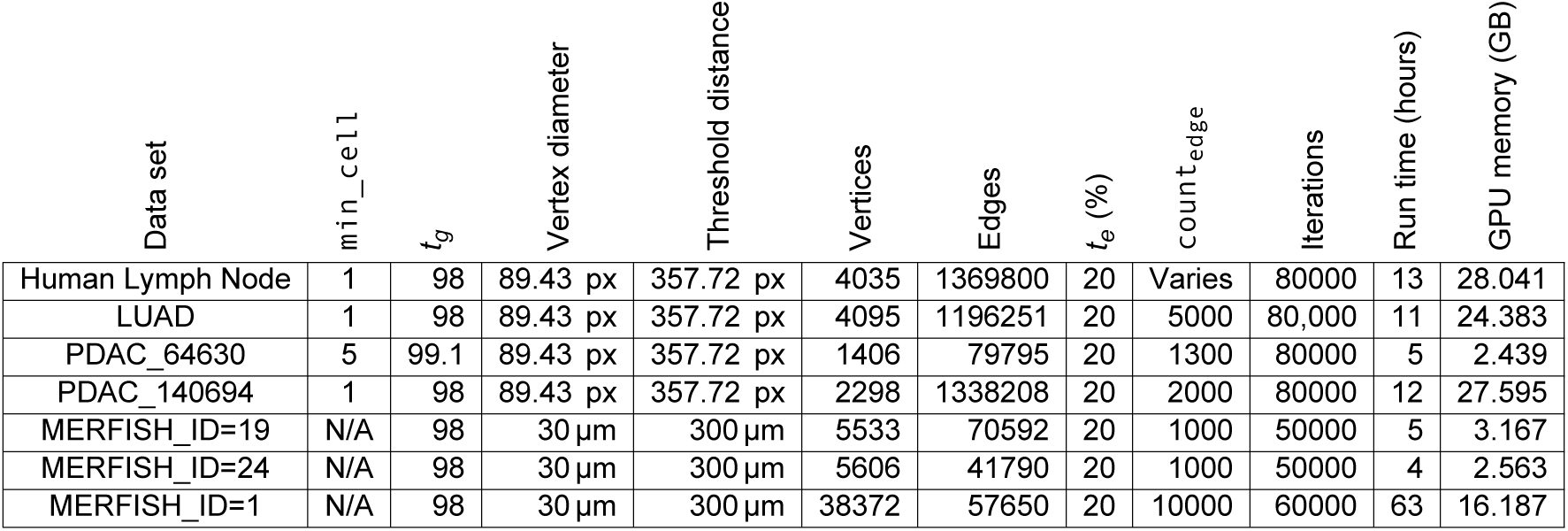
Hyperparameters used for each data set. `min_cell` value is the minimum number of cells used to filter the genes. *t_g_* is the threshold gene expression (in percentile) used for determining active ligand or receptor genes. Vertex diameter represents the spot diameter in pixels for Visium data and cell diameter in micrometers for MERFISH data. Threshold distance is the neighborhood cutoff distance and vertices and edges count the number of each in the input graph. *t_e_* represents the percentage of top CCC we selected for visualization. `count`_edge_ is the threshold for the number of edges with the highest attention scores. Iterations is the number of iterations used to train the model. Each data set also has measured run time and GPU memory.

| Data set | $\min\_cell$ | $t_g$ | Vertex diameter | Threshold distance | Vertices | Edges | $t_e$ (%) | $count_{edge}$ | Iterations | Run time (hours) | GPU memory (GB) |
| --- | --- | --- | --- | --- | --- | --- | --- | --- | --- | --- | --- |
| Human Lymph Node | 1 | 98 | 89.43 px | 357.72 px | 4035 | 1369800 | 20 | Varies | 80000 | 13 | 28.041 |
| LUAD | 1 | 98 | 89.43 px | 357.72 px | 4095 | 1196251 | 20 | 5000 | 80,000 | 11 | 24.383 |
| PDAC_64630 | 5 | 99.1 | 89.43 px | 357.72 px | 1406 | 79795 | 20 | 1300 | 80000 | 5 | 2.439 |
| PDAC_140694 | 1 | 98 | 89.43 px | 357.72 px | 2298 | 1338208 | 20 | 2000 | 80000 | 12 | 27.595 |
| MERFISH_ID=19 | N/A | 98 | 30 $\mu m$ | 300 $\mu m$ | 5533 | 70592 | 20 | 1000 | 50000 | 5 | 3.167 |
| MERFISH_ID=24 | N/A | 98 | 30 $\mu m$ | 300 $\mu m$ | 5606 | 41790 | 20 | 1000 | 50000 | 4 | 2.563 |
| MERFISH_ID=1 | N/A | 98 | 30 $\mu m$ | 300 $\mu m$ | 38372 | 57650 | 20 | 10000 | 60000 | 63 | 16.187 |

### Run time and memory usage

NEST requires a graphical processing unit (GPU), as the program’s core detection module relies on deep learning models. Running time and memory usage depend on the input data size. NEST on the PDAC_64630 sample with 79,795 edges (each representing a relation through ligand-receptor pair) and 1,406 vertices (each representing a Visium spot), took 5 hours with 2.44 GB memory for each run. While we ran most samples with standard 32 GB GPU memory, some (e.g. MERFISH samples) required less memory due to the number of genes (Table 1).

There may be situations where the input data is too large to fit in the available GPU memory. In this case, we offer two options. One option is to split the input graph into multiple subgraphs, for example 4 or 8 subgraphs with overlapping regions, and run each subgraph separately to ensure lower memory consumption (Supplementary Figure S5a). When benchmarking the split input graph against the total input graph, we observed similar performance for both synthetic and pancreatic cancer samples (Supplementary Figure S5b versus Figure 3c, and Supplementary Figure S5c-d versus Supplementary Figure S5e-f). If the split input graph is infeasible, we apply a filtering option to reduce the number of genes based on the parameter `min_cell` or raise the threshold gene expression percentile *t_g_* so fewer genes will be activated. Increasing the value of `min-cell` will reduce the number of genes in the dataset, thereby reducing the input graph size, which has a negligible impact on the outcome (Supplementary Figure S6). These options for handling extensive datasets enable NEST’s usage on a wide variety of machines and increasingly large molecular information.

### Synthetic data preparation for benchmarks

We generated 3,000 equidistant data points represeting Visium spots, each having 10,000 genes. We assigned 10% of genes as ligand or receptor genes and formed synthetic ligand-receptor pairs with these genes. The synthetic ligand-receptor database generated this way has ≈ 1,400 pairs. In this same way, we sampled 5,000 data points from a uniform distribution representing MERFISH cells, each having 350 genes. The synthetic ligand-receptor database generated this way has 100 pairs with 12% of genes acting as ligand or receptor genes to approximate observed proportions^19^. Lastly, we sampled 5,000 data points from a mixture of uniform and Gaussian distribution representing single-cell data types, each having 350 genes, with 12% genes forming ligand-receptor pairs. The synthetic ligand-receptor database generated this way has 100 pairs.

### Threshold distance setup for neighborhood cutoff

Our threshold distance for neighborhood formation is 89.34 px × 4 for Visium data, where each spot diameter is 89.34 This threshold forms a neighborhood (direct communication) within 3 hops. For MERFISH samples, we set the threshold distance to be 100×3 µm, where the cell diameter is approximated as 100 µm for 3 hops. Users may change these parameters in the Data Preprocessing Step using `–spot_diameter` and `–neighborhood_-threshold` hyperparameters.

### Spatial transcriptomics of human pancreatic cancer patients

Two solid tumor biospecimens were collected from the pancreas of two stage IIB PDAC patients (PDAC_64630: 76 year-old male, PDAC_140694: 83-year old female). Both biospecimens were collected from the University Heath Network Biospecimens Program (Toronto, Canada). Ethical approval was obtained through the University Health Network Research Ethics Board (13-6377). Tumors were collected at the time of resection. Samples were stabilized for approximately 3 hours at 4 °C until long term preservation (embedded in optimal-cutting-temperature compound). Samples were stored at −80 °C until used. The cases were selected according to have >30% tumor celullarity. The regions of interest for capture areas (6.5 mm ×6.5 mm) were selected, targeting tumor areas with representative subtype morphologies^57^. 10 µm cuts were placed into 10x Genomics Visium FFPE spatial gene expression slides from selected trimmed tissue areas. Spatial transcriptomics using the Visium platform was carried out according to manufacturer’s instructions (10X Genomics. Part number: 1000200, Protocol : CG000160 RevB, CG000239 RevD). Sequencing was performed on the Illumina NovaSeq 6000 platform with paired-end reads according to 10x Genomics specifications. Data was processed using Space Ranger (version 2.0.0) and mapped to the GRCH38 v93 genome assembly.

### Annotation of pancreatic cancer samples

Histology categories of tumor and stroma were assigned based on the following features — tumor: malignant cells arranged any architecture of glands, cords, strands, solid sheets and single cells^57^; stroma: non-tumor tissue surrounding tumor cells, composed mainly of fibroblasts, myofibroblasts, and collagen fibers^56^. Transcriptomic subtype annotations were assigned using Loupe Browser version 6.4.1 (10X Genomics) according to Log2 Feature Sum filter using a previously determined subtype gene list^64^.

### PDAC patient-derived organoid library

An organoid library with matching whole transcriptome sequencing from laser microcapture enriched tumors was established from 44 cases with resectable (stage I/II) and advanced (stage III/IV) pancreatic ductal adenocarcinoma. Tumor transcriptomic subtype classification was obtained from published data^55^. Advanced organoids were generated by University Health Network Living Biobank as part of a clinical trial (no. NCT02750657) and resectable organoids were generated at the Notta Lab (CAPCR 13-6377, 21-5648) following established methods^82^. After passage 6, RNA was extracted from dissociated organoids. Sequencing libraries were prepared using the Smart-3SEQ protocol^83^ from 10 ng of RNA. Pools of 20 libraries were sequenced on the Illumina NextSeq 500 using 150 cycles kit v2 for Single Read 150 on a Mid Output flow cell.

### Development of the interactive NEST

NEST Interactive uses vanilla Javascript and HTML on the front-end with Tailwind CSS for styling. We used D3.js for the histogram and Vasco Asturiano’s 3d-force-graph library (which extends off of D3.js and Three.js) for the responsive graph. To obtain the data for display, NEST Interactive uses jQuery to send AJAX requests to the back-end server as well as to deep-copy current graph data. The back-end uses the Django framework. After receiving a request from the front-end with edge count as a parameter, a Python script reads all CSV records stored locally and returns graphable nodes and edges in JSON format. Necessary files to be read include complete records for cell (or spot), cell coordinates, cell annotations (if available), and the list of top 20% CCC detected by NEST. NEST Interactive further processes this data by separating vertices into connected components and assigning colours using NumPy, Pandas, SciPy, and Matplotlib libraries. NEST Interactive is available at: https://github.com/schwartzlab-methods/NEST-interactive.

## Data availability

PDAC spatial transcriptomic data for PDAC_64630 and PDAC_140694 are available on the Gene Expression Omnibus under accession number to be provided before publication. The spot annotations for both samples are available at https://github.com/schwartzlab-methods/NEST_paper_figures/blob/main/NEST_figures_input_PDAC.7z. We obtained spatial transcriptomic data of human lymph node from https://www.10xgenomics.com/datasets/human-lymph-node-1-standard-1-0-0, mouse hypothalamic preoptic region from https://datadryad.org/stash/dataset/doi:10.5061/dryad.8t8s248, and LUAD from the Gene Expression Omnibus under accession number GSE189487.

## Code availability

NEST is available at https://github.com/schwartzlab-methods/NEST with a tutorial at https://github.com/schwartzlab-methods/NEST#vignette. Scripts for generating the figures and plots of the manuscript can be found at https://github.com/schwartzlab-methods/NEST_paper_figures.

## Acknowledgments

We thank Melanie Peralta from the Pathology Research Program Laboratory at the University Health Network, for her work on the processing and mounting of pancreatic resections into Visium slides. This work was supported by the Canadian Cancer Society Challenge Grant (grant 707484; G. W. S.), the Natural Sciences and Engineering Research Council of Canada (grants RGPIN-2023-04713 and DGECR-2023-00395; G. W. S.), the Social Sciences and Humanities Research Council (grant NFRFE-2022-00681; G. W. S.), the Canada Research Chairs Program (G. W. S.), the Princess Margaret Cancer Foundation (G. W. S.), the Gattuso-Slaight Personalized Cancer Medicine Fund & Research Stimulus Grant 2022 from the Princess Margaret Cancer Foundation (F. N.), Ontario Institute for Cancer Research (F. N.), the Ontario Early Researcher Award (grant ER19-15-205; F. N.), and the University of Toronto’s Eric and Wendy Schmidt AI in Science Postdoctoral Fellowship, a program of Schmidt Futures.

## Author contributions

G. W. S. conceived and supervised the project. F. T. Z. developed the NEST method, software, and benchmarks. J. L. developed the Nest Interactive software. F. T. Z. ran and analyzed benchmarks. E. F. generated experimental results. F. T. Z., E. F., and D. P. ran and analyzed data. F. T. Z., E. F., J. L., D. P., and G. W. S. wrote and edited the manuscript. All authors reviewed the manuscript.

## Supplementary Information

### Supplementary Figures

**Supplementary Figure S1:**
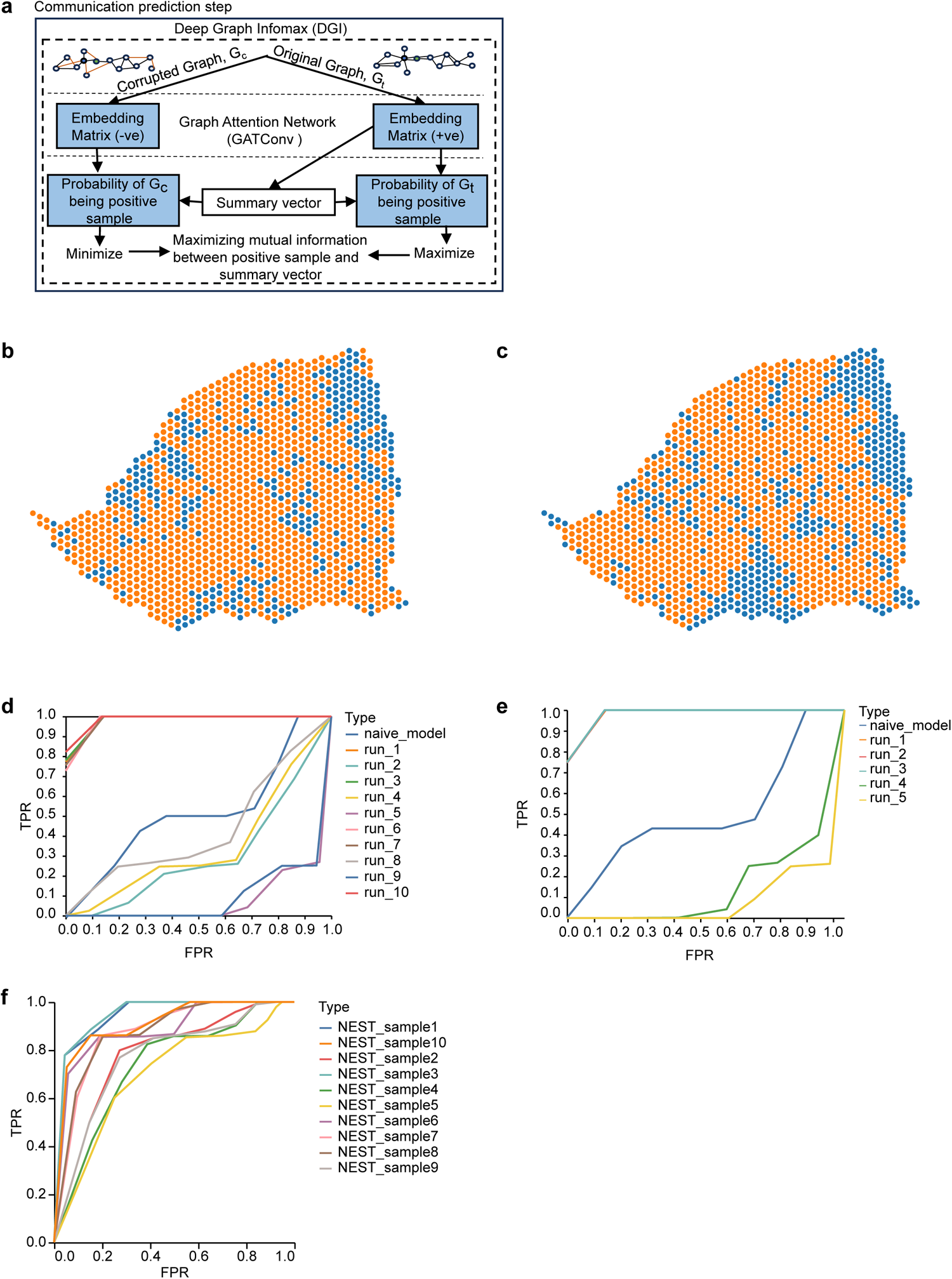
Schematic overview of the cell-cell Communication Prediction Step using the DGI framework (**a**; see Methods) and additional benchmarking illustrating naïve model behavior and instability of alternative activation functions (**b-f**). **a** DGI is a contrastive learning framework. We used a graph attention network as the encoder in this framework. The workflow consists of two branches: the corrupted branch and original branch. The corrupted branch applies random permutations to the edges of the original graph *G_t_* to obtain a corrupted graph *G_c_*, or negative sample, as input to an encoder, generating a negative embedding. The original branch uses *G_t_* as the positive sample, obtaining the positive embedding using the same encoder as the corrupted branch. DGI uses the positive embedding to produce a summary vector representing the global representation of *G_t_*. DGI maximizes the mutual information between the summary vector and the positive embedding, and maximizes the Jensen-Shannon divergence between the summary vector and the negative embedding. This process returns the trainable parameters learned using DGI. **b-c** PDAC_64630 Visium data with spots colored by high (top 5% scoring, orange) and low (blue) input ligand-receptor co-expression scores (**b**) or high (top 5% scoring, orange) and low (blue) CCC prediction scores using a naïve model (**c**). The naïve model’s output mostly overlaps input ligand-receptor co-expression scores, suggesting a lack of spatial integration and ligand-receptor pattern measurement. **d-e** ROC curves of NEST with PReLU (**d**) and ReLU (**e**) activation functions for attention calculations across multiple runs. These curves demonstrate unstable performance compared to the main NEST model with Tanh activation. **f** ROC curves for NEST performance on ten random synthetic data sets generated from uniform distributions show the increased stability of NEST across variable data sets and model initialization.

**Supplementary Figure S2:**
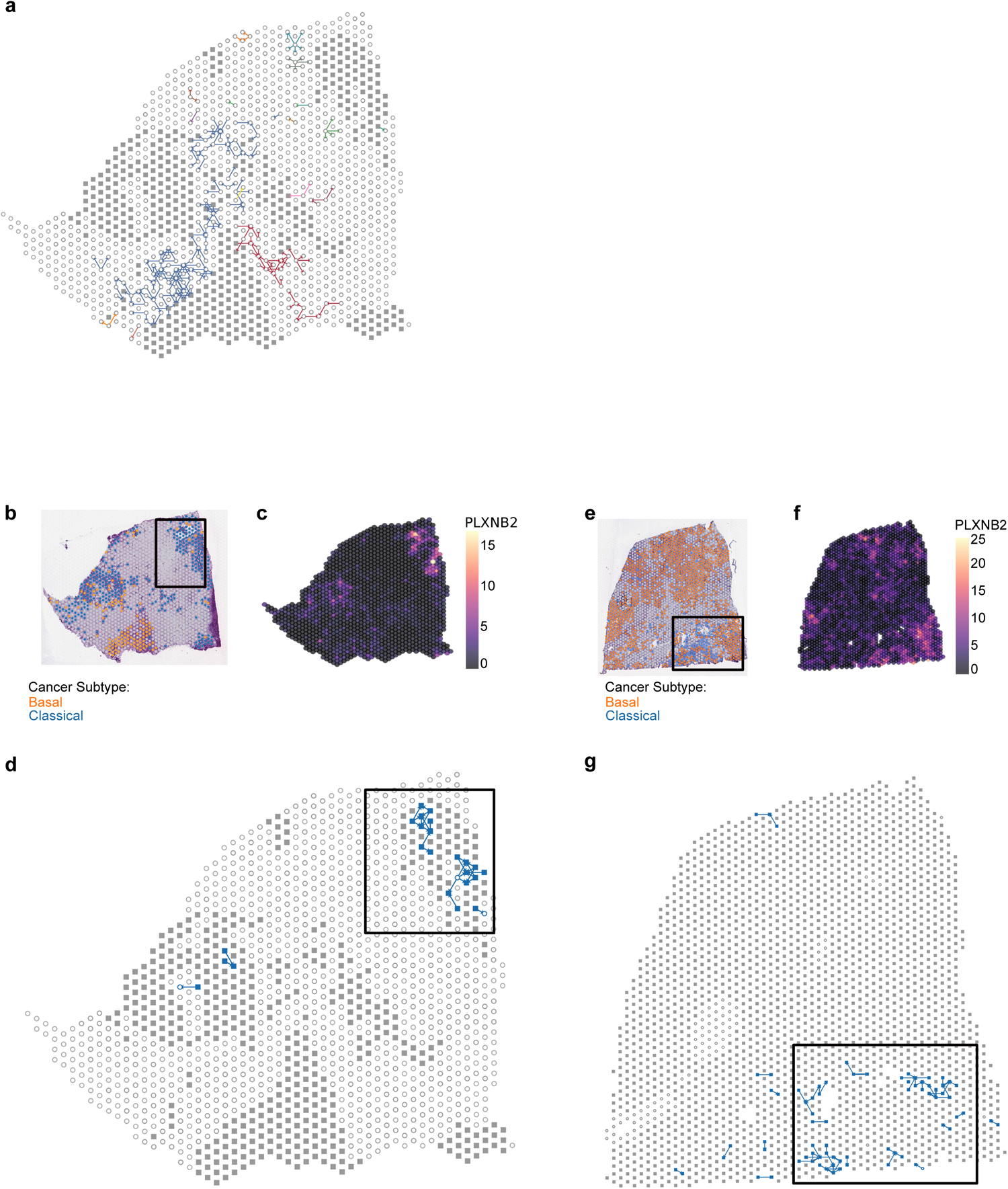
Enrichment of specific CCC in Classical regions of PDAC tissue. **a** NEST output of PDAC_64630 with communication filtered for *FN1*-*RPSA*. Colors represent connected components from Figure 6c with tumor (filled square) and stromal (open circle) spots. *FN1*-*RPSA* is found mostly within the stromal region. PDAC_64630 Visium data with spots colored by subtype (**b**), *PLXNB2* expression (**c**), or connected component from the NEST output filtered for *PLXNB2* signals (**d**). NEST associates *PLXNB2*-initiated communication in Classical regions, highlighted with a black box. **e-g** As with b-d but in PDAC_140694.

**Supplementary Figure S3:**
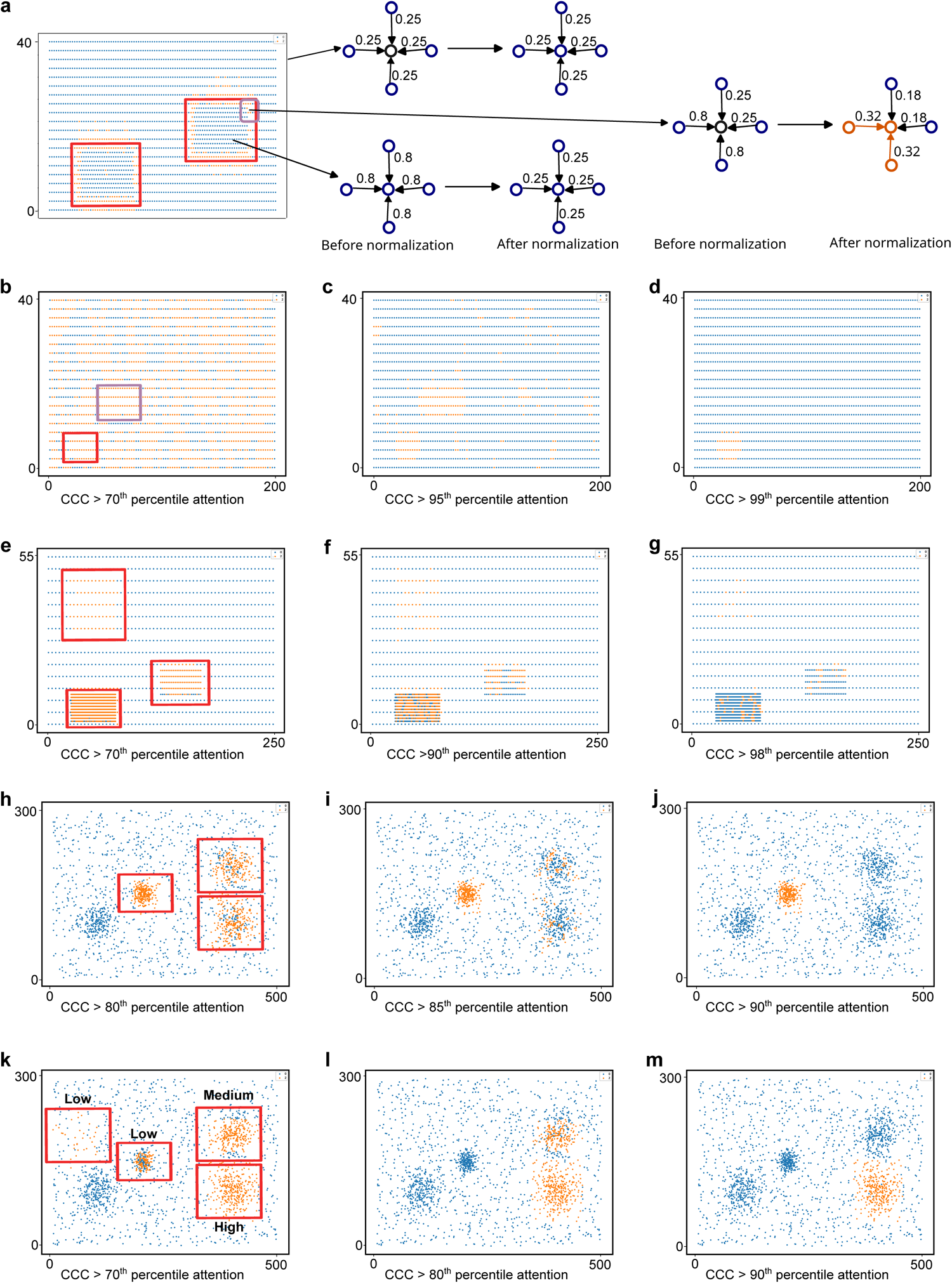
NEST generally assigns higher attention scores to ligand-receptor pairs with higher input ligand-receptor co-expression scores. Points represent cells or spots colored by no communication (blue) or detected communication with at least one other point (orange). Red boxes on images within the left column highlight *a priori* assigned active regions with higher input ligand-receptor co-expression scores. **a-d** Synthetic data sets with uniform, equidistant distribution. **a** Detected CCC with normalized attention scores per vertex (see Methods). Vertices near boundaries, both within and outside of the assigned high co-expression score region, were detected as the strongest CCC. These vertices had both low attention and high attention edges, so after normalization, the high attention edges were increased. In the center of the high co-expression score region, there was an even split of high attention edges, so normalizing edges within a vertex lowered attention scores. This phenomenon illustrates the need for unnormalized attention scores for global comparison. **b-d** Example output of NEST applied to a data set with two high co-expression score regions with increasing percentile expression of 70 (**b**), 95 (**c**), and 99 (**d**). With unnormalized scores, correct regions are identified. The purple box disappears first (**d**) as it contains lower ligand-receptor co-expression scores as compared to the red box. **e-g** NEST output for increasing percentiles applied to data with high co-expression score regions also with high density. **h-j** NEST applied to data with a mixture of uniform and Gaussian distribution (varying pairwise distance between the cells) with a same level of ligand-receptor pair co-expression. NEST resolves the race between pairwise distance and ligand-receptor co-expression between cells and assigns higher attention scores to higher density regions (**j**). **k-m** NEST applied to data with a mixture of uniform and Gaussian distribution with different levels of ligand-receptor pair co-expression. NEST recovers high co-expression score regions even with lower density than other, lower activity regions.

**Supplementary Figure S4:**
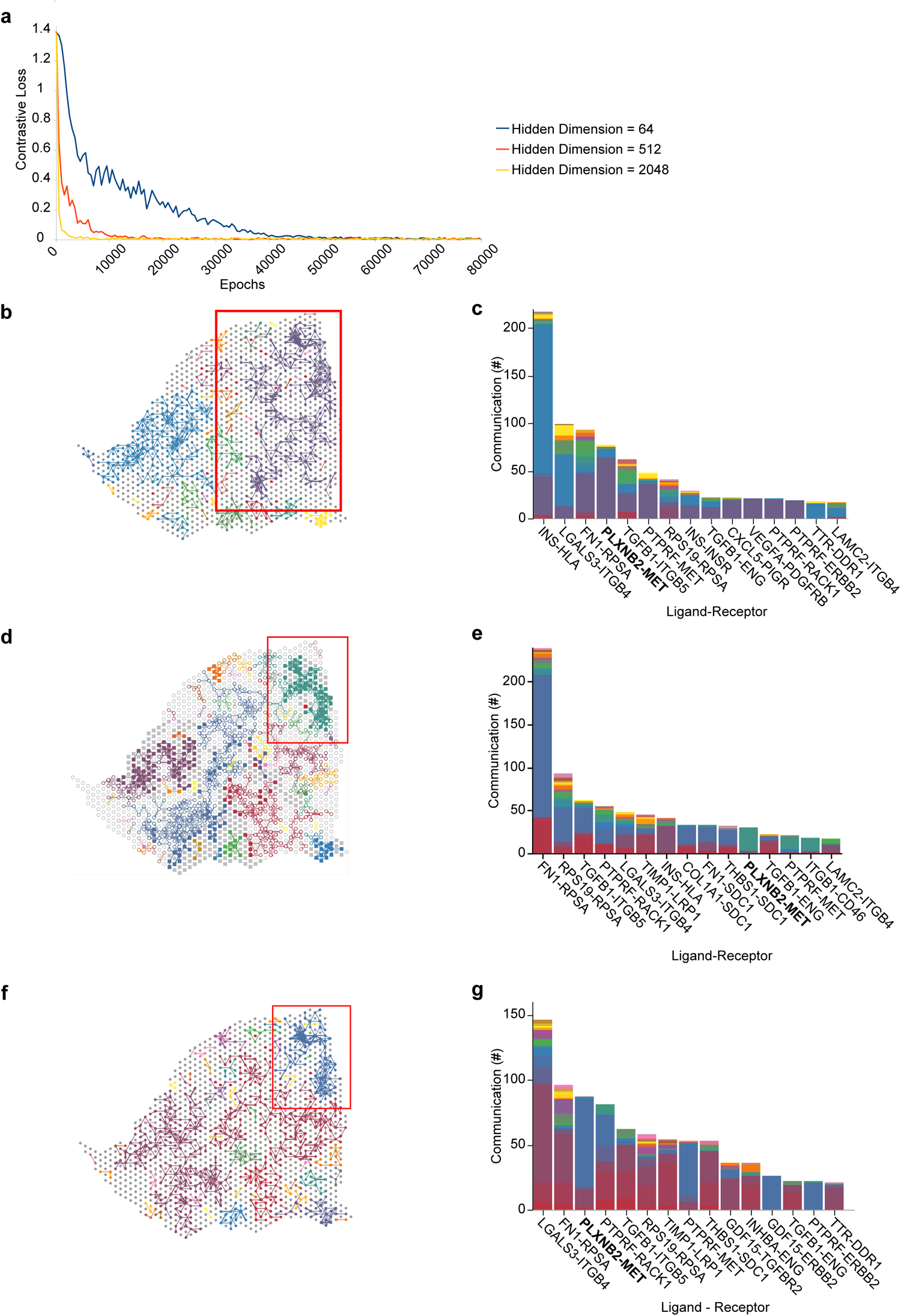
Larger hidden dimension size improves model training. **a** Loss (*x*-axis) vs. iterations (*y*-axis) curve of NEST for three hidden dimension sizes (64, 512, and 2048) on PDAC_64630. **b-g** Connected component plots (**b, d, f**) for PDAC_64630 generated by NEST with 64 (**b**), 512 (**d**), and 2048 (**f**) hidden layer dimensions paired with corresponding histograms (**c, e, g**) of the most abundant ligand-receptor pairs from **b**, **d**, and **f**, respectively. Higher dimensions (e.g. 512 and 2048) are good at localizing communication by providing more disjoint components associated with exclusive ligand-receptor pairs as shown by red boxes on the connected component plots (e.g. *PLXNB2*-*MET*, highlighted with bold text). However, to control GPU memory consumption we chose 512 as the hidden dimension size to balance model performance as well as computational complexity and memory.

**Supplementary Figure S5:**
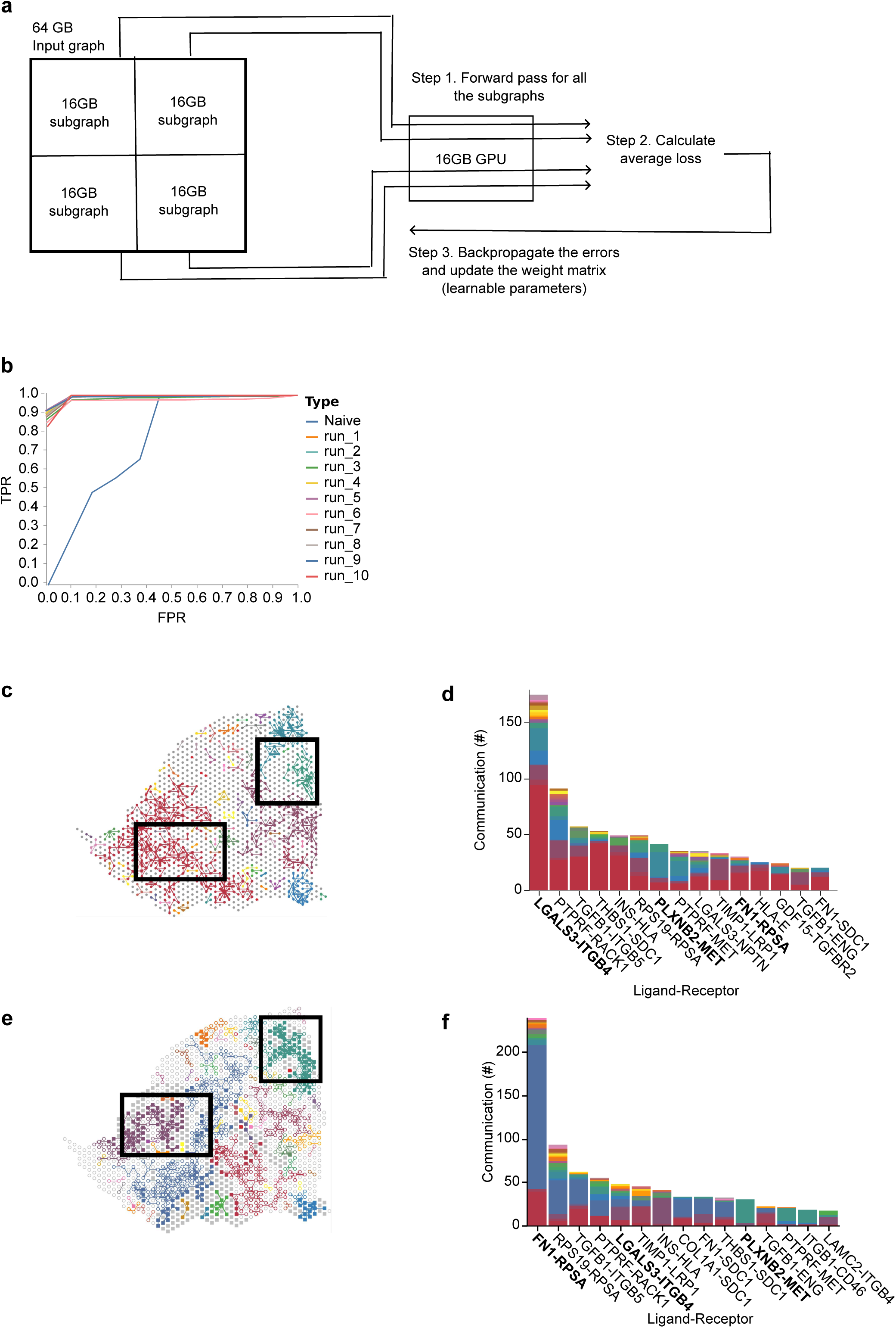
Splitting the input graph into multiple overlapping subgraphs for separate processing reduces GPU memory consumption. **a** Flowchart showing the training procedure for split input. For example here, input data needs 64GB GPU memory, whereas only 16 GB GPUs are available. We split the input graph into four subgraphs (overlapping is not shown just for simplicity) each requiring 16GB GPU memory. We pass subgraphs through the GPU sequentially (one 16 GB GPU available) or in parallel (multiple 16GB GPUs available). After one round of the forward pass (Step 1), we gather the loss for all subgraphs and take the average loss (Step 2) to backpropagate in order to update the learnable weight parameters (Step 3). These three steps run in a cycle for a specified number of iterations or until the model has converged. After model convergence, we use the learned attention scores as previously described to predict the CCC at different locations (or subgraphs) of the input. **b** ROC curves from 10 runs of NEST for synthetic data (equidistant data points) with split-input case. The split-input algorithm (lower memory consumption) shows equally good performance as the default non-split input case as provided in Figure 3c-e of the main manuscript (higher memory consumption). **c-d** NEST output visualization (**c**) colored by connected component on PDAC_64630 using split input with corresponding histogram of the most abundant ligand-receptor pairs (**d**). **e-f** As with **c-d** but with non-split input. With and without split input have high overlap for both the CCC network visualization (highlighted with black box) and the most abundant ligand-receptor pairs (bold text).

**Supplementary Figure S6:**
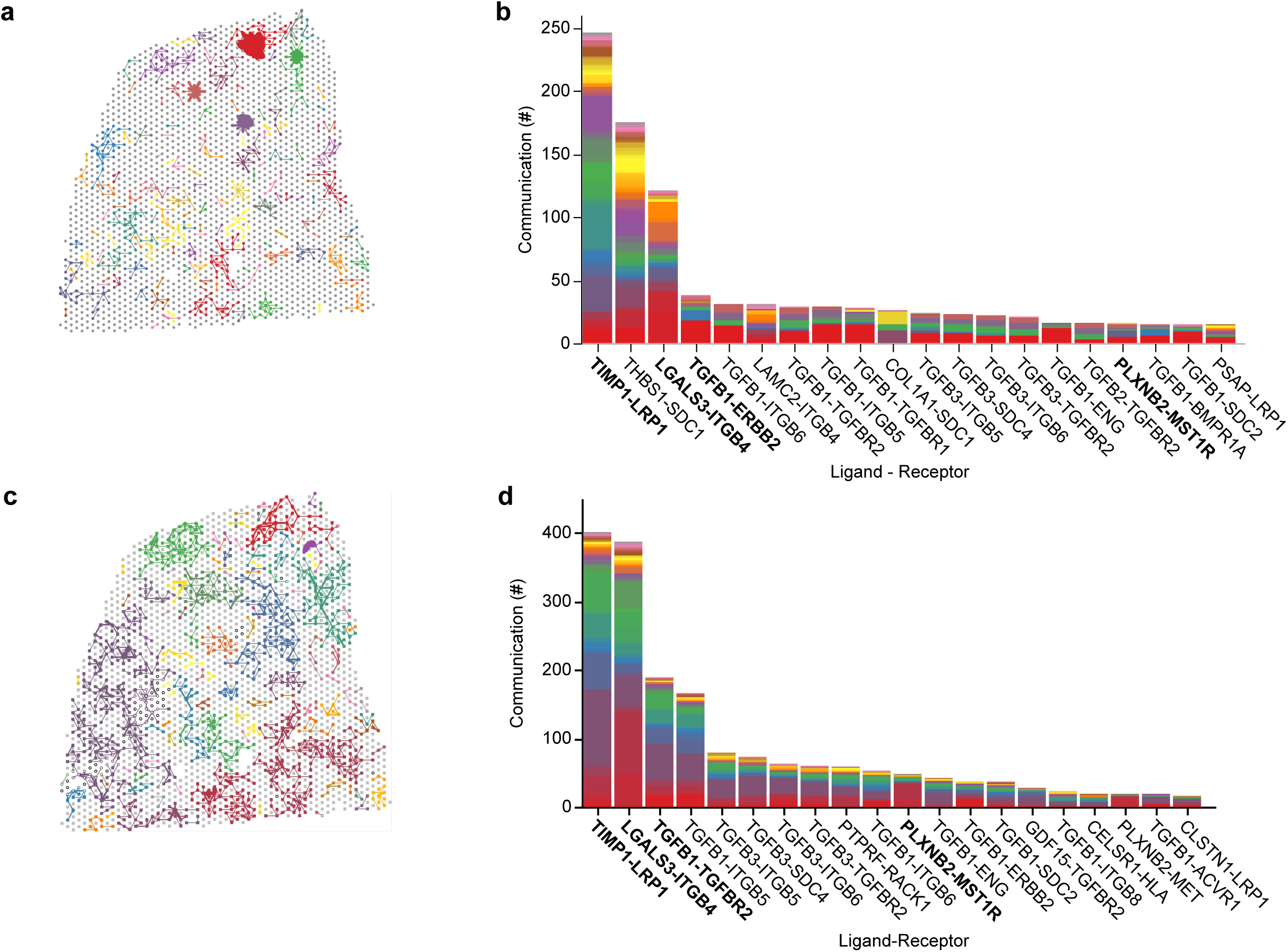
Impact of gene filtering to reduce the input data size for lower memory consumption. **a-b** CCC visualization with colored connected components (**a**) and corresponding histogram of the most abundant ligant-receptor pairs (**b**) from NEST applied to PDAC_140694 with `min_cell` = 10 results in less than one million edges in the input graph. **c-d** As with a-b but with default NEST. The default NEST applied to PDAC_140694 contains over one million edges in the input graph using the default preprocessing setup with `min_cell` = 1. Both inputs generate similar ligand-receptor pairs as shown with bold text in the histograms. Therefore, we use higher `min_cell` value to reduce GPU memory consumption if necessary.

**Supplementary Figure S7:**
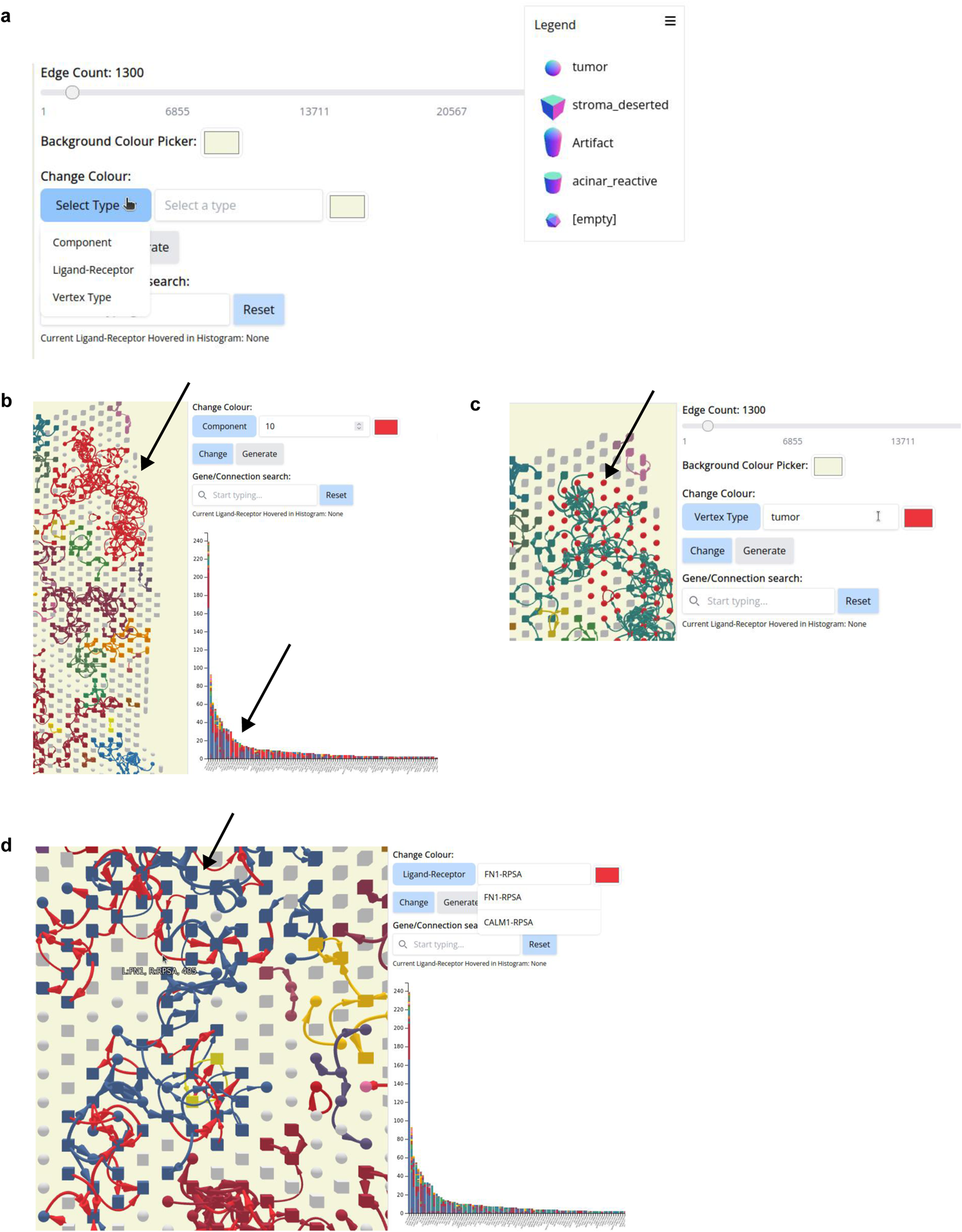
NEST Interactive is a feature-rich visualization for data exploration of spatial communication. **a** User input fields, including color pickers for components, ligand-receptor pairs and vertex types. **b-d** Example of changing colors of connected component 10 (arrow; **b**), tumor vertex type (**c**), or ligand-receptor *FN1*-*RPSA* (**d**) to red.

**Supplementary Figure S8:**
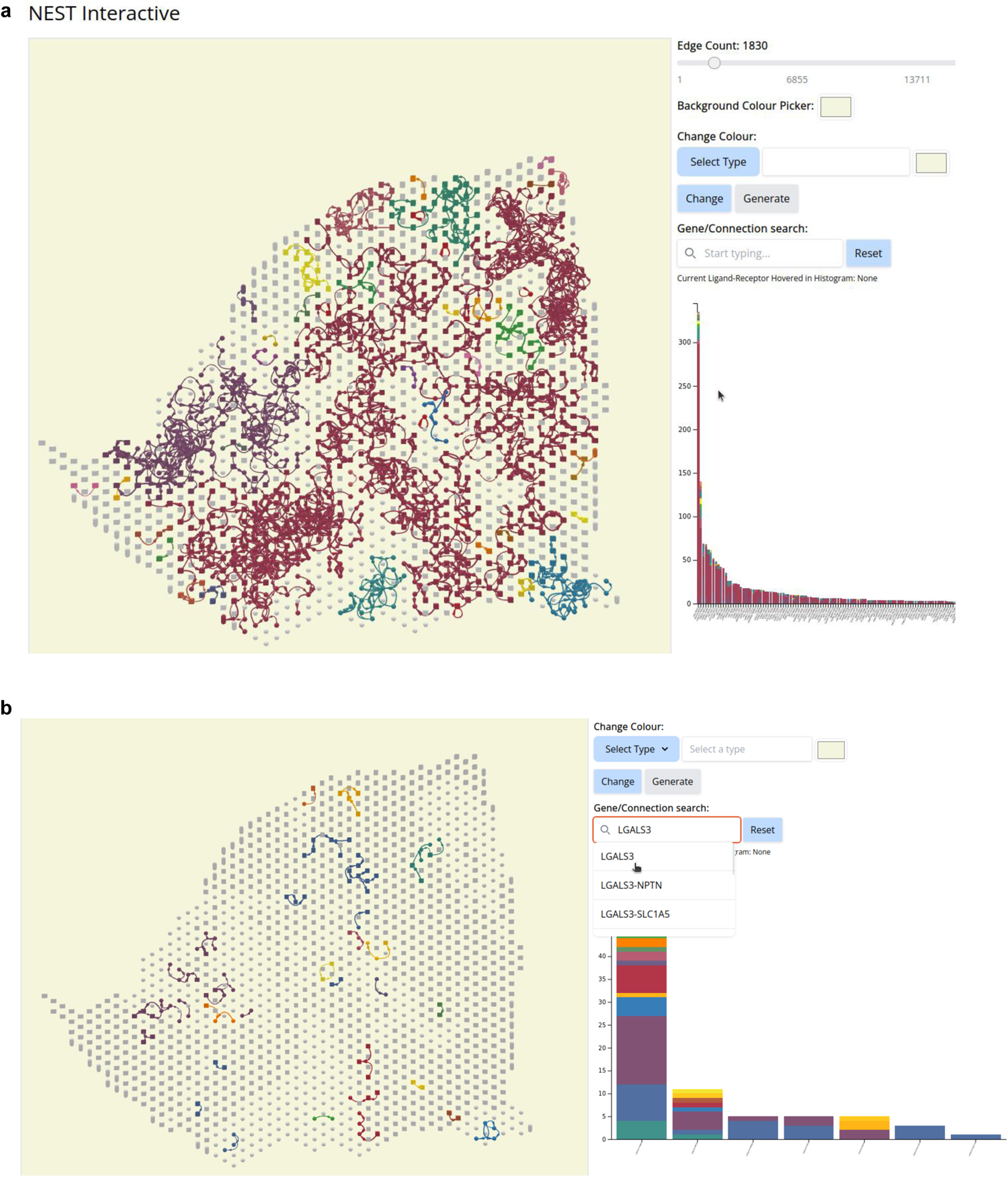
NEST Interactive simplifies filtering for specific ligand-receptor pairs. **a** The sliding Edge Count bar enables the user to specify the number of top communicating edges to visualize. Here, we select the top 1830 CCC. **b** The gene or connection search option allows the user to focus on a particular ligand-receptor pair only. This field implements a fuzzy search for the users to choose from a list of available genes and pairs.

### Supplementary Notes

#### Supplementary Note S1: Chi-squared and hypergeometric test for an enrichment analysis of *PLXNB2*-*MET* in the Classical region of PDAC tissue

To determine ligand-receptor pair usage in the Classical region (connected component 10 of PDAC_64630; top right green region in Figure 6c), we performed an enrichment analysis of *PLXNB2*-*MET*. We applied a chi-squared test on the histograms generated from connected component 10 to determine whether there was a bias in ligand-receptor pairs occurrence. We found that was indeed a skew in the 57 ligand-receptor pair types (Chi-squared test: *p* = 6.14 × 10^−32^, Χ^2^ = 282.34, degrees of freedom = 56), suggesting enrichment of certain types of CCC in connected component 10. Of these skewed types, we found *PLXNB2*-*MET* communication is enriched in the Classical region (Hypergeometric test: *p* = 1.41 × 10^−22^).

We performed the enrichment analyses using `scipy.stats.chisquare` and `scipy.stats.multivariate_hypergeom` from SciPy. Scripts for this analysis are available at https://github.com/schwartzlab-methods/NEST_paper_figures/.

#### Supplementary Note S2: Statistical analyses of PDAC patient-derived organoid model bulk RNA-seq data

To compare gene expression among bulk RNA-seq counts of PDAC patient-derived organoids, we normalized counts with the DESeq2^84^, followed by log2 transformation, and z-score normalization (Figure 7a).

To compare the expression of specific genes, we normalized bulk RNA-seq counts of organoids with DESeq2 (Figure 7b-c). We found one Basal organoid model as an outlier and excluded from analysis upon further investigation due to an atypical *MYC* amplification. We compared the expression of *MET* (Fisher-Pitman permutation test: *p* = 3.18 × 10^−2^ and *LGALS3* (Fisher-Pitman permutation test: *p* = 0.175) between Classical (*n* = 5) and Basal (*n* = 5) organoids using the exact two-sample Fisher-Pitman permutation test from the `oneway_test` function from the `Coin` package (Figure 7b-c). Scripts for performing each analysis are available at https://github.com/schwartzlab-methods/NEST_paper_figures/.

